# Genetic modulation of immune gene co-expression in the aged mouse hippocampus by the *Apbb1ip* locus

**DOI:** 10.1101/2025.06.04.657838

**Authors:** James E. Tomkins, Aditi Methi, Gerard Bouland, Birte Doludda, Kitty Murphy, Jeremy A. Miller, Rupert W. Overall, Khyobeni Mozhui

## Abstract

Ageing is a major risk factor for many neurodegenerative diseases, and the hippocampus is particularly vulnerable to the effects of ageing. To define the transcriptomic changes related to ageing and the impact of genetic variation, we analysed hippocampal gene expression data generated from a genetically diverse panel of inbred mouse strains belonging to the BXD Family. We applied a combination of differential expression, differential correlation, and weighted gene co-expression network analyses, followed by genetic mapping to delineate age-associated transcriptomic patterns and genetic modulators. This study revealed an upregulation in immune response and microglial genes. We identified a key age-associated co-expression module enriched in immune related genes, and quantitative trait locus mapping of this module uncovered a genetic regulatory locus in which *Apbb1ip* was the primary candidate gene. Additionally, gene level differential correlation analysis identified a substantial restructuring of the hippocampal transcriptome during ageing. Notably, *Ywhab,* an important signalling chaperone displayed altered gene expression correlations with >70 genes, between young and aged mice. These results provide novel insights into transcriptional dynamics of hippocampal ageing and identified the immune gene *Apbb1ip* as a potential modulator of immune response and microglial gene upregulation with implications for neurodegenerative disease pathogenesis.

**Highlights:**

- Complementary gene expression analyses indicated an upregulation of immune responses in the ageing hippocampus of a genetically diverse mouse cohort, likely driven by microglia subtypes.
- A systems genetics approach revealed the *Apbb1ip* locus as a modulator of immune gene co-expression, highlighting a novel candidate regulator of age-associated neuroimmune dynamics.
- Signalling chaperone *Ywhab* gene expression was highly differentially correlated with other transcripts during ageing which may underlie widespread age-associated molecular perturbations in the hippocampus.

## Introduction

Ageing is a primary risk factor for numerous diseases, including many neurodegenerative diseases^1,2^. As global demographics shift toward an older population, the healthcare, economic and social burdens due to age-related diseases of the central nervous system are expected to increase dramatically^3^. Despite research investment, our understanding of molecular mechanisms underlying neurodegenerative disease pathogenesis remains incomplete. This has hampered therapeutic development, and people diagnosed with these diseases currently have limited treatment options^4,5^.

One promising approach to address these challenges is to characterise the defining features of the ageing brain. Cellular hallmarks include synaptic dysfunction, mitochondrial impairment and inflammation^6^. Moreover, different brain regions exhibit differential vulnerability to the ageing process^7^. The hippocampus, a central player for cognitive performance, including executive function and working memory, is particularly vulnerable to volumetric changes and age-related neuroinflammation (inflammageing)^8,9^.

Underlying cellular changes of brain ageing are molecular perturbations. Transcriptomics analyses are a powerful approach to uncover global gene expression patterns at a gene level resolution and infer implications for molecular pathways. Gene expression studies have identified a plethora of processes that contribute to brain ageing. These include decreased synaptic function, increased immune activation, differential splicing, and dysregulated genome maintenance^10^. Interestingly, transcriptomics studies of brain ageing have highlighted gene expression signatures similar to those observed in Alzheimer’s disease, particularly in the hippocampus^11^.

Recent advances in RNA sequencing technologies, at single-cell/nucleus and spatial resolutions, have specifically implicated glial involvement and immune activation in the ageing brain^12,13^. Additionally, bulk RNA sequencing of the ageing mouse hippocampus has identified age-dependent changes related to synaptic function and immune activity^14,15,16^. However, it is important to note a critical limitation of transcriptomics studies of the ageing mouse brain, that these were almost all performed using a single mouse strain, C57BL/6J, often regarded as the representative laboratory mouse. Restricting analyses to a single genetic background impedes the generalisability to other mouse strains, let alone to other species^17,18,19^.

In order to incorporate genetic diversity in characterising transcriptomic signatures of the ageing hippocampus, we utilised a publicly available microarray dataset of hippocampal gene expression in a cohort of BXD mice^20,21^. BXDs are a population of recombinant inbred mice originally derived from crosses of the C57BL/6J and DBA/2J strains^22,20^. Collectively, this genetic reference panel harbours significant genetic variation with approximately 6 million common genetic variants segregating across the different strains. Due to this, the BXD Family exhibits substantial phenotypic variation, including traits related to ageing and lifespan. As a relevant model for cognitive ageing, members of the BXDs show significant differential vulnerability to age-related cognitive function^23,24,25^. The BXDs therefore serve as a powerful systems genetics platform for mapping genetic loci that influence gene expression dynamics related to ageing and age-related disease.

Using this resource, our goals were to identify age-related gene expression networks, evaluate their dynamic restructuring with ageing and assess how common genetic variants modulate this process. Specifically, we performed differential expression analysis (DEA) and weighted gene co-expression network analysis (WGCNA) to detect linear expression changes with ageing, and quantitative trait locus (QTL) mapping to interrogate genetic modulation. Additionally, we applied differential correlation analysis (DCA) to gain new perspectives on how gene-gene correlations are altered over the life course.

Our results confirm that changes in immune and microglial expression signatures are a salient feature of the ageing hippocampus^26,9,27^. Importantly, the QTL mapping identified a regulatory region on chromosome 2 where the immune activation gene, *Apbb1ip*, is a strong candidate modulator of an age-associated immune co-expression module. Furthermore, the DCA revealed extensive restructuring of gene-gene associations during ageing and highlighted signalling chaperone *Ywhab* as a highly differentially correlated transcript (differentially correlated with 71 other genes) between young and aged animals.

## Methods

### Data Acquisition and Pre-processing

This study utilised microarray-based gene expression data available on the GeneNetwork web platform^28^ (http://www.genenetwork.org/), derived using the GeneChip™ Mouse Gene 1.0 ST Array (ThermoFisher #901168), a whole-transcriptome array consisting of 35,558 transcript-level probes. The dataset consists of hippocampal gene expression from 234 mice belonging to the BXD genetic reference population^20^ (GeneNetwork accession: GN712; GEO accession: GSE274733). Data had been normalised using the RMA (Robust Multiarray Average) algorithm, log-transformed and standardised to a mean of 8 units with standard deviation of 2 units. The processed dataset was obtained as a flat file from GeneNetwork.

Probes that were not annotated to coding transcripts were excluded, resulting in a total of 26,273 gene expression traits in 234 samples (sample information in **Data S1**). For further sample level quality assessment, we used two strong Mendelian gene expression traits that are *cis*-regulated and map as expression QTL in *cis* (*cis*-eQTLs): *Zfp125* on chromosome (Chr) 12 (probe ID 10399588), and *Mkl2* on Chr16 (probe ID 10433656). We leveraged these strong Mendelian traits as genotype proxies to impute strains and verified consistency between reported and imputed strains. Based on this, we detected two samples of uncertain genotypes (BXD65b_1 and BXD75_1) that did not match the imputed genotypes, and these were excluded. For outlier detection, we performed hierarchical clustering using the *hclust* function in R (method = “average”). This is part of initial data pre-processing for WGCNA (see below). This clustering analysis identified 9 outlier samples, and these were also excluded. All downstream analyses were performed using 223 samples (analysis cohort) that were retained after the sample level quality assessment. Details of the analysis cohort sample age and sex demographics are provided in **Table 1** and **Data S1**, including sample categorisation into Young/Mid/Aged age groups.

**Table 1:** Sample age and sex demographics of analysis cohort. Following initial quality assessment of the dataset these samples were used for downstream analysis.

| Age (days) | Age Category | N<sub>samples</sub> | Sex |  |
| --- | --- | --- | --- | --- |
|  |  |  | Male | Female |
| < 480 | Young | 82 | 36 | 46 |
| 480 - 600 | Mid | 70 | 29 | 41 |
| > 600 | Aged | 71 | 29 | 42 |

### Differential Expression Analysis

Differential expression analysis (DEA) was performed in R using a linear model, where gene expression was the outcome variable and age in days was the predictor variable (continuous), with sex and batch as cofactors. The Benjamini-Hochberg procedure was used to correct for false discoveries and significance was assumed at P_FDR_ ≤ 0.05 (**Supplementary Appendix 1)**. Differentially expressed genes (DEGs) were subjected to functional enrichment analysis using Reactome annotations and the g:Profiler g:SCS calculation^29^, the background gene set for determining enrichment was the collection of genes on the array.

### Cell Type Enrichment Tests

Expression weighted cell type enrichment (EWCE)^30^ was used to identify cell types underlying age-related gene expression changes in the mouse hippocampus. EWCE requires 3 inputs: (i) a target gene list; (ii) a background gene list; (iii) a gene-by-cell-type specificity matrix. Using these, EWCE determines the probability that the target genes have higher expression specificity in each cell type than would be expected by chance, achieved through bootstrapping enrichment tests. Here, our target gene list was DEGs in the ageing mouse hippocampus, derived from our DEA. The background gene list included all genes expressed in the mouse hippocampus. The gene-by-cell-type specificity matrix was generated using *Tabula Muris Senis*^31^, a whole-body single-cell transcriptome dataset of the ageing mouse (ageing time points 3, 12 and 18 months), and the *generate_celltype_data* function in the EWCE package. Given that we were specifically interested in the hippocampus, data was subset to include only brain cell types. Following recommendation from the EWCE developers, we conducted 10,000 bootstrap enrichment tests, p-values were FDR corrected and significance assumed at P_FDR_ ≤ 0.05.

### Weighted Gene Correlation Network Analysis

We constructed gene co-expression networks using the weighted gene co-expression network analysis (WGCNA) package^32^ (version 1.70.3) in R (**Supplementary Figure 2A**). The normalised expression values were used to compute pairwise Pearson correlations between transcripts and to define a co-expression similarity matrix. A soft-thresholding power of 10 was used to transform the similarity matrix into an adjacency matrix (R^2^ = 0.88, mean connectivity = 136; **Supplementary Figure 2B**). Next, the pairwise topological overlap matrix (TOM) and the corresponding dissimilarity values (1-TOM) between genes were calculated to build a signed gene co-expression network. Co-expression modules were defined using the following parameters: minimum module size of 30 using the *blockwiseModules* function with specified arguments [networkType = “signed”, TOMtype = “signed”, deepSplit = 3, pamStage = T, pamRespectsDendro = F, reassignThreshold = 0], otherwise default parameters. Modules that were closely related were merged using a 0.2 dissimilarity correlation threshold. This resulted in 33 modules (**Figure 2A**) and the module eigengenes (MEs) were then used to test association with age, based on the assigned age categories. The R code is provided in **Supplementary Appendix 1**.

### Gene Overlap Test, Pathway Analysis and Network Visualisation

The R package GeneOverlap (version 1.26.0) was used to test the intersection of gene lists using Fisher’s exact test^33^. Gene expression signatures of mouse hippocampal cell types were defined based on the “Mouse Whole Cortex and Hippocampus 10x” dataset from the Allen Brain Map^34^. This dataset includes >1 million cells collected from the hippocampus and cortex of 8-week-old male and female mice. We defined hippocampal cell types as any cluster from which >25% of cells were collected in a hippocampal dissection. For glial types without any clusters meeting this criterion, we selected the relevant cluster with the highest fraction of hippocampal cells. Clusters were then assigned to broader types based on subclass annotation, totalling 12 defined cell types: oligodendrocyte (OLG), microglia/perivascular macrophage (MG/PVM), oligodendrocyte precursor cell (OPC), CA1 excitatory neuron (Ex-CA1), CA2 excitatory neuron (Ex-CA2), CA3 excitatory neuron (Ex-CA3), interneuron (InN), dentate gyrus excitatory neuron (DG), endothelial cell (END), smooth muscle cell/pericyte (SMC/Peri), astrocyte (AST) and vascular and leptomeningeal cells (VLMC). OPCs were not explicitly annotated but cluster 365_Oligo was assigned as OPCs based on expression of *Olig1*, *Olig2*, and *Pdgfra* (**Supplementary Figure 3**). Marker genes were defined by sorting genes based on the expression level (log_2_(CPM+1)) in one cell type minus the maximum expression in the other 11 types. The top 50 genes were used for gene overlap analysis, whereby these marker gene lists were intersected with co-expression module gene lists. Functional annotation and enrichment analysis of gene lists were performed using the clusterProfiler R package (version 3.18.1)^35^, unless otherwise specified. Network visualisations were created using Cytoscape (version 3.9.1)^36^.

### Quantitative Trait Locus Analysis using GeneNetwork

All MEs and gene expression traits are publicly available on GeneNetwork^28^. The MEs can be directly accessed from the website using the following query: Species: Mouse; Group: BXD Aged Hippocampus; Type: Traits and Cofactor; Dataset: B6D2RI Phenotypes; and by using the search term “WGCNA”. For expression traits, Type: Hippocampus mRNA; and Dataset: UTHSC BXD Aged Hippocampus Affy Mouse Gene 1.0 ST (Jun15 RMA), and transcripts can be retrieved by gene symbol. Quantitative trait locus (QTL) mapping was performed using the Genome-wide Efficient Mixed Model (GEMMA) option^37,38^. All genome coordinates for QTL and gene locations are based on the GRCm38/mm10 mouse reference genome.

### Differential Correlation Analysis

To test differences in correlation coefficients of gene expression for pairs of genes or co-expressed modules between age groups, differential correlation analysis (DCA) was performed, whereby the young and aged mice were selected for the contrast (see **Table 1** & **Data S1**). Differential correlation was determined similarly as previously described^39^. Spearman’s rank correlations between pairs of genes were separately calculated in the two groups (young and aged) resulting in two correlation coefficients for each pair of genes. Next, the correlation coefficients were transformed to z-scores using Fisher z-transformation^40^. As z-scores are inherently normally distributed, they are amenable to a two-sample z-test, resulting in a Δz-score representing the probability of whether the change in correlation coefficient is based on chance. The variances required for the two-sample z-test were calculated by 1.06/(n-3), with n being the sample size of the respective groups. As Δz is normally distributed, a two-sided P-value for the differential correlation between each pair of genes was calculated. The Benjamini-Hochberg procedure was used to correct for false discoveries and significance was assumed at P_FDR_ ≤ 0.05. Differential correlations were classified as: gain of association (strengthening of the correlation coefficient between young and aged cohorts); loss of association (weakening of the correlation coefficient between young and aged cohorts); or flip of association (when the correlation coefficients are within 0.1 between young and aged cohorts, but in the opposite direction, i.e., a change from positive to negative correlation or vice versa). In addition to pairwise gene correlations, this analysis approach was also applied to MEs derived from the WGCNA.

### Protein Interaction Analysis

Protein interaction prediction and assessment of reported physical protein-protein interactions or associations were performed for 14-3-3-beta/alpha (gene name *Ywhab*). For prediction of 14-3-3 binding motifs within the corresponding proteins which were differentially correlated with *Ywhab* (at the transcript-level) between young and aged mice, the 14-3-3-Pred tool was utilised^41^. This protein sequence-based resource adopts three machine learning (ML) classifiers (artificial neural networks, position-specific scoring matrices and support vector machines) to evaluate the binding likelihood of 14-3-3 proteins, based on prior ML model training. The list of differentially correlated transcripts was queried and scored in a binary manner whether each of the ML classifiers deemed a motif in the protein of interest a positive hit, based on confidence thresholds predefined by the tool developers^41^. This resulted in a prediction score ranging from 0 to 3 for each protein.

Whether any of the differentially correlated gene protein products were previously reported to interact with 14-3-3-beta/alpha was assessed using the Molecular Interaction Search Tool (MIST; v5.0)^42^. Ywhab was queried for mouse protein-protein interactions and low rank results were filtered out. Gene/protein synonyms were considered, and these were then assessed in the context of the differentially correlated transcripts.

## Results

### Description of samples

The analysis cohort consisted of hippocampal transcript expression data for 223 mice of the BXD Family (129 females and 94 males). Age ranged from 223 to 972 days. Strain, sex, and age demographics for each sample are provided in **Data S1**. There was no significant difference in age between the sexes (mean age of 550 ± 112 days for females, and 539 ± 113 for males). After grouping samples by age into young, mid and aged categories, there were 82, 70 and 71 samples per age group, respectively (**Table 1**).

### Immune-related genes are upregulated with age

To investigate age-associated differential gene expression, we applied a linear regression model to the 26,273 transcripts, with sex and batch as covariates (full results provided in **Data S2**). At an FDR of 5%, only 37 transcripts were significantly differentially expressed with age (**Figure 1A**). Thirty-four of these were upregulated, and three transcripts were downregulated with age. Expression of immune related *C4b* (complement C4B, involved in complement activation), and *Ccl3* (C-C motif chemokine 3, a proinflammatory factor) were the most robustly upregulated genes with age, based on p-value: both P_FDR_ < 0.001. *Npbwr1* (Neuropeptides B/W receptor type 1) was the most downregulated gene with age, P_FDR_ = 0.029. The 37 differentially expressed genes (DEGs) were significantly enriched for 5 Reactome terms related to immune function (**Figure 1B**). The most significant term was “Innate Immune System” (P = 2.98 x 10^-3^).

**Figure 1:**
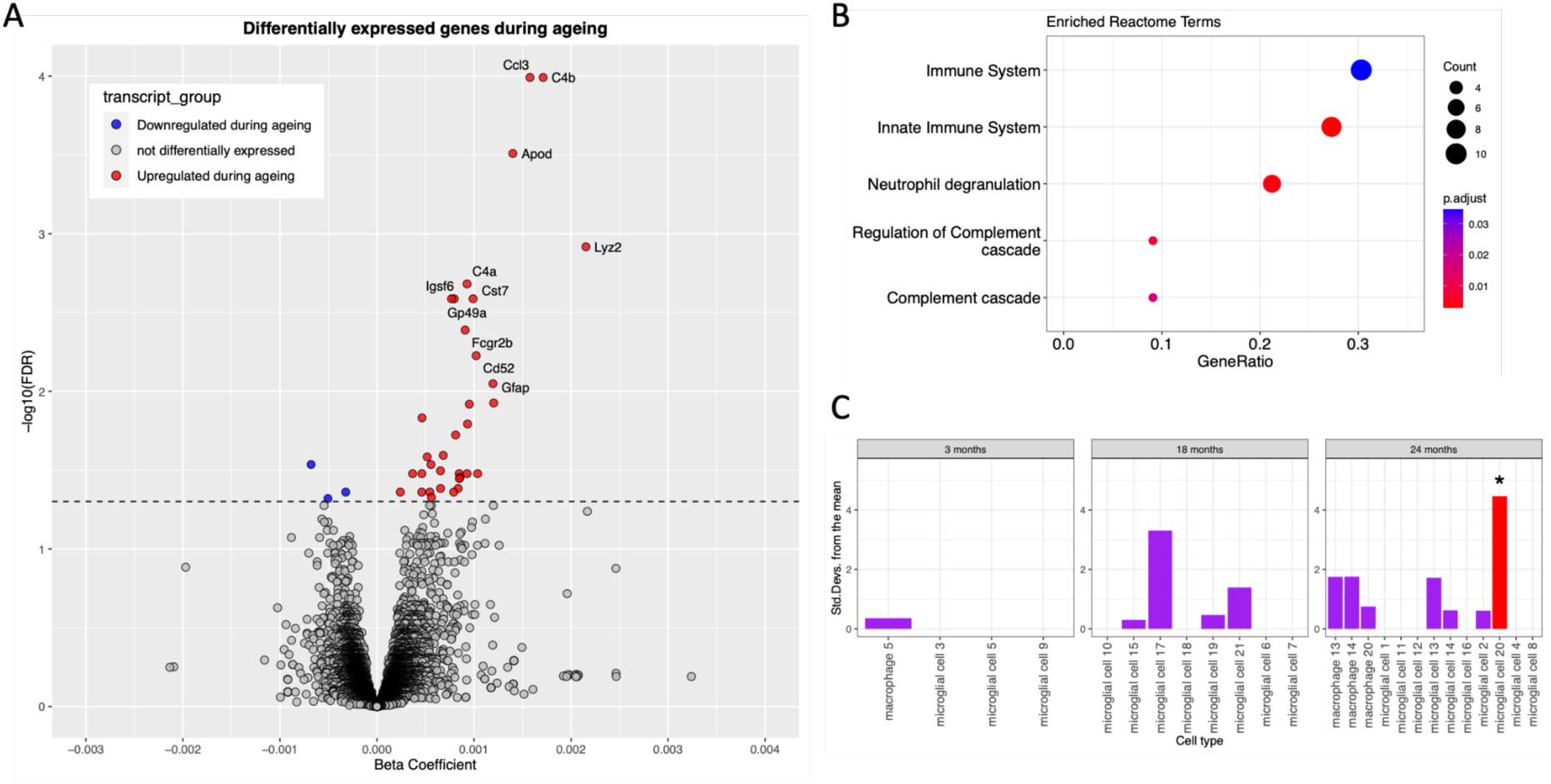
Hippocampal transcriptome differential expression analysis across age in BXD mice. **(A)** Volcano plot of age-dependent differentially expressed transcripts, calculated with age (in days) as a continuous variable. Statistical significance threshold defined at 5% false discovery rate (FDR). **(B)** Functional enrichment analysis of differentially expressed transcripts using Reactome annotations. GeneRatio represents the proportion of transcripts in the query list (n=33 Reactome annotated transcripts) assigned the corresponding enriched Reactome term, similarly represented as absolute count by dot size. **(C)** Cell type enrichments calculated using Expression Weighted Cell type Enrichment (EWCE)^30^. EWCE was performed using DEGs from the ageing mouse hippocampus and a subset ageing mouse whole body single-cell dataset specifically for brain cell types of myeloid origin^31^. Microglial cell 20, a microglia cluster specific to 24-month-old mice, was exclusively enriched for the DEGs after FDR correction.

**Figure 2:**
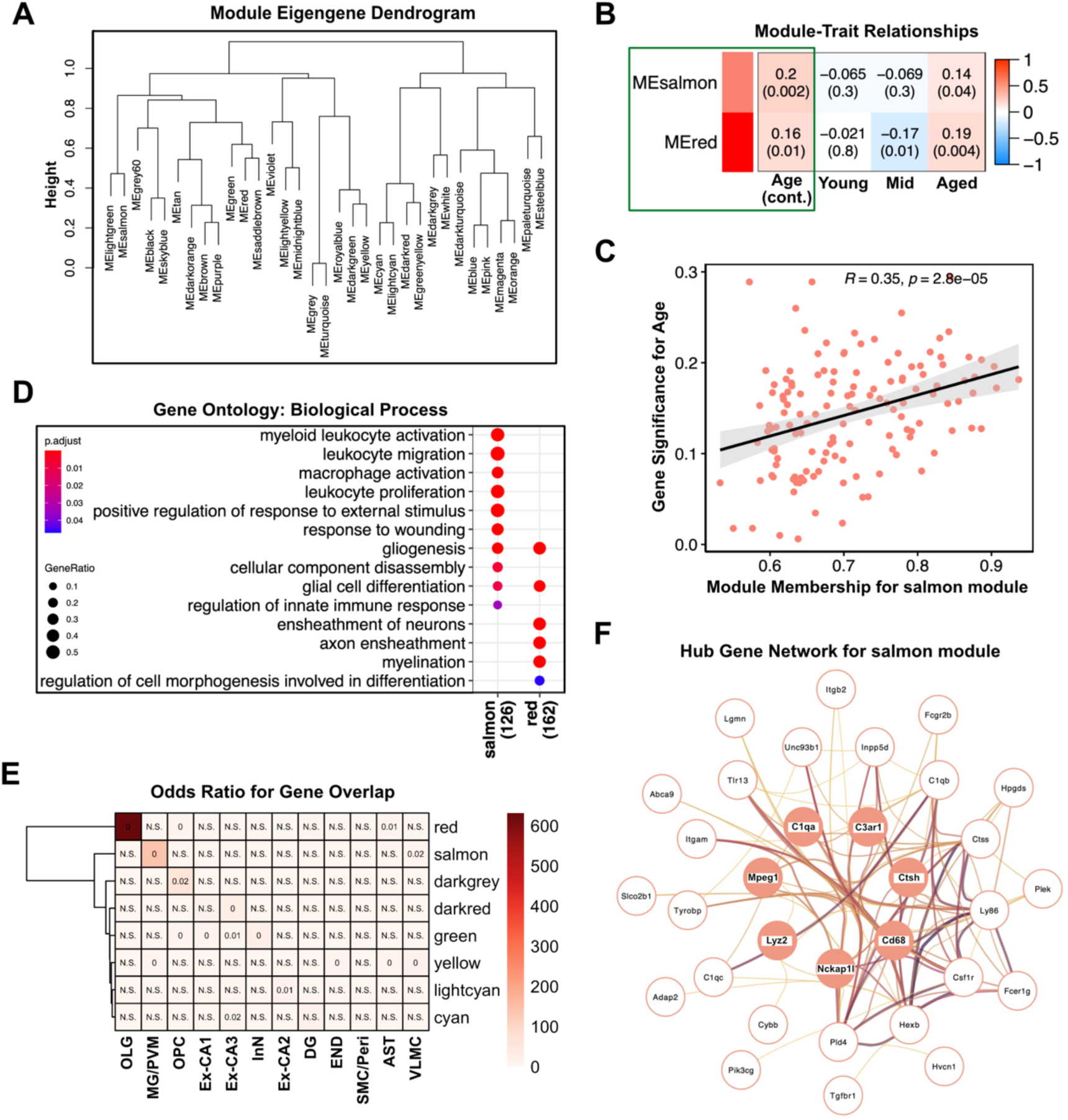
Weighted gene co-expression network analysis of ageing hippocampal samples from BXD mice revealed co-expressed immune response genes that significantly correlate with age. **(A)** Hierarchical clustering dendrogram of module eigengenes for 33 co-expressed gene modules. **(B)** Module-trait heatmap showing two modules (“salmon” and “red”) with statistically significant correlation between their module eigengene and the age variable. The intensity of the colour indicates the extent of module correlation. Positive correlation is denoted by red, negative correlation by blue. The correlation coefficient and p-value (in brackets) for each module-trait relationship is displayed. **(C)** Pearson correlation between module membership values and gene significance (correlation of gene expression profile with age) of all genes in the salmon module (R = correlation coefficient and p = p-value). **(D)** Gene ontology biological process functional enrichments for co-expressed genes in the salmon and red modules. The numbers in brackets along the x-axis denote the total number of module genes that were annotated and used in the analysis. **(E)** Heatmap showing overlap (calculated using Fisher’s exact test) between genes in selected modules and markers for major cell types in the brain (OLG = oligodendrocytes, MG = microglia, PVM = perivascular macrophages, OPC = oligodendrocyte precursor cells, Ex-CA1 = CA1 excitatory neurons, Ex-CA2 = CA2 excitatory neurons, Ex-CA3 = CA3 excitatory neurons, InN = interneurons, DG = dentate gyrus excitatory neurons, END = endothelial cells, SMC = smooth muscle cells, Peri = pericytes, AST = astrocytes, VLMC = vascular and leptomeningeal cells). The intensity of the colour indicates the magnitude of the odds ratio representing the strength of association between the two gene lists. The text within the heatmap denotes the p-value for each association. **(F)** Hub-gene network for salmon module, showing hub genes at the centre and their first-and second-degree neighbours in subsequent shells. The nodes were selected to have a module membership of at least 0.5 and gene significance (for age) of at least 0.15. The hub genes were selected based on a threshold of 0.8 for module membership and 0.2 for gene significance. Edge weights denote the adjacency which were filtered to retain adjacency > 0.05.

We hypothesised that the upregulation of immune-related genes detected from bulk expression data was likely driven by immune cell types of the brain. To investigate this, we tested which brain cell type markers are overrepresented in the set of identified age-related DEGs, using Expression Weighted Cell Type Enrichment (EWCE)^30^. This method utilises single-cell transcriptomic profiles to identify “driver” cell types based on the expression specificity of a given set of genes. Here, we used the 37 DEGs alongside brain expression profiles from *Tabula Muris Senis*, a whole-body single-cell transcriptome dataset of the ageing mouse^31^. A EWCE analysis on brain cell types of the myeloid lineage revealed an enrichment for a microglial cluster specific to 24-month-old mice (**Figure 1C**), suggesting that among myeloid cells of the brain the most extreme differential expression changes as observed in bulk DEA can be explained by a specific subtype or cell state of microglia in the aged brain. A follow-up analysis using an independent single-cell dataset encompassing a broad range of cell types in the mouse brain^43^ highlighted a consistent enrichment in microglia for genes differentially expressed in our DEA (**Supplementary Figure 1A**). In addition, a strong association with perivascular macrophages was also evident. Interestingly, 4 DEGs from our DEA were also markers of the recently described age-dependent microglia (ADEM) subtype^44^ (**Supplementary Figure 1B**). Furthermore, the ADEM marker genes were overrepresented in the same age associated microglia cluster identified by EWCE of DEGs in the *Tabula Muris Senis* dataset (**Supplementary Figure 1C & Figure 1C**). Taken together, the DEA performed on bulk expression data coupled with EWCE leveraging single-cell transcriptomics revealed molecular and cellular responses which support neuroimmune upregulation in the ageing hippocampus.

### Immune response and myelination co-expression modules are associated with hippocampal ageing

To complement DEA, we used a network approach to define groups of tightly co-expressed genes related to hippocampal ageing. We performed WGCNA^32^ on the 26,273 transcripts, across 223 samples. At a soft thresholding power of 10 (**Supplementary Figure 2B**), the topological overlap matrix-based clustering resulted in 33 co-expression modules that ranged in size from 32 transcript members for the violet (smallest) module, to 4396 transcript members for the turquoise (largest) module (module description in **Data S3** and transcript-level module membership is specified in **Data S2**). Each module represents a network of co-expressed transcripts, and the predominant variation of each module is captured by its module eigengene (ME), defined by the first principal component of the module (ME values and proportion of variance explained by each ME are provided in **Data S1 & S3**). Hierarchical clustering of MEs indicated that many of the modules are strongly intercorrelated (**Figure 2A**). Next, we characterised these modules by examining their associations with age and sex. For age, we found positive correlations with the ME of the salmon (*r* = 0.2, *p* = 0.002) and red (*r* = 0.16, *p* = 0.01) modules (**Figure 2B**). The salmon and red modules consist of 136 and 186 transcript members, respectively. This relationship implies that the overall expression of genes in these modules increases with age.

To gain biological insight into the functional representation of these modules (and others), we performed Gene Ontology (GO) enrichment analyses for all modules, separately (**Data S4**). 18 of the 33 modules showed significant enrichment (*P*_FDR_ < 0.05) for specific Gene Ontology Biological Processes (GO BPs), and we classified each of these by their most enriched functional categories. For example, the red module was classified as “myelin related” based on the top enriched GO BP terms “ensheathment of neurons”, “axon ensheathment” and “myelination” **(Figure 2D)**. The primary functional category of each module is listed in **Data S3**. Furthermore, we identified several modules that have modest associations with sex and may represent sex-specific expression networks in the ageing hippocampus, such as dark orange (glycolytic processes), salmon (immune response) and purple (mitochondrial protein complex) modules (**Data S3**).

The salmon (immune response) module was selected for further analysis as it had the strongest association with age (**Figure 2B**). Each transcript member of a module is assigned a module membership value (computed as the correlation between the expression of that gene and the ME). For the salmon module, there was a significant positive correlation between gene module membership and correlation with age, demonstrating that genes most associated with age also tend to be the most central elements of this age-associated module (**Figure 2C**). Similar to the age associated DEGs, of which 35% were components of the salmon module, the genes in the salmon module were significantly enriched for proinflammatory immune processes, such as leukocyte and macrophage activation (**Figure 2D**). Members of the salmon module also showed a significant overlap with marker genes of microglia and perivascular macrophages (**Figure 2E**), further indicating a role of immune cells in the ageing hippocampus. Notably, the red (myelin related) module genes were very strongly associated with oligodendrocyte cell markers.

A network of the most highly connected genes in the salmon module (that also showed significant association with age) was constructed to visualise the key co-expression interactions between hub genes and their first-and second-degree connections (**Figure 2F**). Interestingly, four of these seven hub genes (*C1qa, C3ar1, Cd68, Lyz2*) were also upregulated during ageing, as observed in the DEA. In summary, additional to the univariate DEA which identified 37 age-associated genes based on individual gene-level tests, our co-expression network analysis uncovered a far broader landscape of age-related inflammatory signals within the hippocampal transcriptome.

### *Apbb1ip* locus modulates age-associated immune response co-expression module

We next sought to investigate the genetic modulation of the salmon (immune response) co-expression module. Considering the salmon ME as a quantitative trait, we performed quantitative trait locus (QTL) mapping and identified a significant module-level expression QTL (module-QTL)^45^ on chromosome (Chr) 2. The strongest QTL linkages were to markers at 18–25 Mb (*p* = 4 x 10^-5^; **Figure 3A**). Among the 136 transcript members of the salmon module, only *Apbb1ip* is located within this peak module-QTL interval, at 22.77 Mb. We then performed an eQTL analysis of the *Apbb1ip* transcript. The hippocampal expression of *Apbb1ip* mapped as a strong *cis*-regulatory eQTL (*cis*-eQTL), indicating that genetic variants within the *Apbb1ip* locus exert a *cis*-regulatory effect on its expression (**Figure 3B**). The peak linkage markers were located on Chr 2 at ∼22 Mb (*p* = 6.9 x 10^-11^) and overlapped the salmon module-QTL. *Apbb1ip* expression is highly specific to microglia and immune cells of the hippocampus (**Supplementary Figure S4**) and is modestly increased in expression with ageing (age effect *p* = 0.04; **Data S2**). The gene with the highest module membership in the salmon module was *Pld4* (5’-3’ exonuclease, located on Chr 12 at 112.76 Mb; **Data S2**). A similar eQTL analysis showed that expression of *Pld4* maps as a modest *trans*-eQTL to *Apbb1ip* (peak markers at Chr 2, ∼23 Mb, *p* = 0.0008; **Figure 3C**). Note that expression of the *Pld4* transcript also maps close to its locus on Chr 12, however, the peak markers are located at ∼84 Mb and hence this is not considered a *cis*-eQTL. Collectively, this genetic analysis indicated that tight co-expression among the salmon module members is not only due to age-related influence, but also partly due to co-regulation by a strong eQTL at the *Apbb1ip* locus.

**Figure 3:**
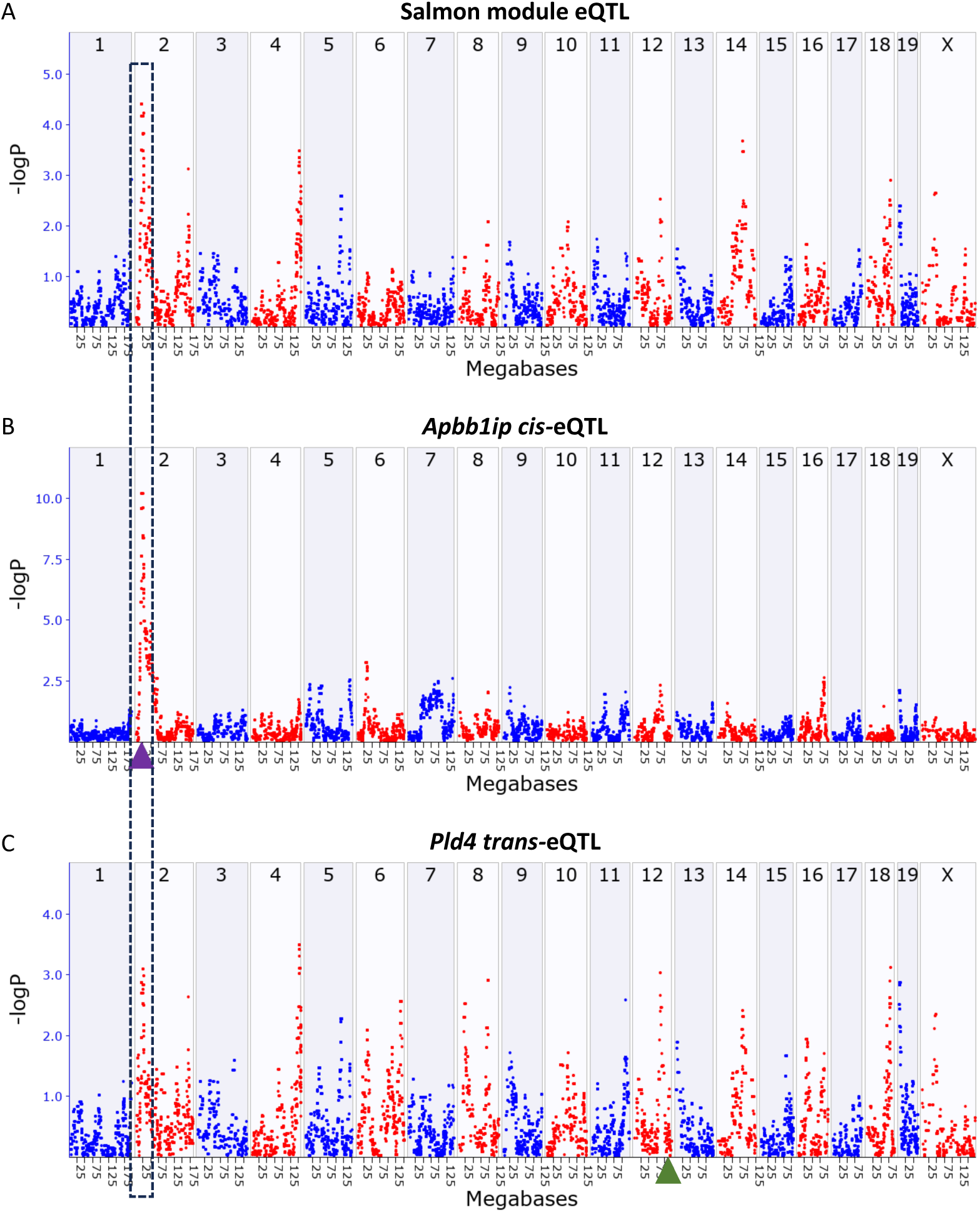
Genetic co-regulation of immune genes by the Apbb1ip locus. **(A)** The salmon module (network of co-expressed immune response genes) genetic regulation landscape represented as a quantitative trait locus (QTL) Manhattan plot. The x-axis indicates the genomic location of each genetic marker, the y-axis represents the linkage-log_10_P. The QTL for the salmon module eigengene has peak linkage on chromosome (Chr) 2 at the Apbb1ip locus, a transcript member of the salmon module. **(B)** Expression of Apbb1ip in the hippocampus maps as a cis-eQTL. The purple triangle (on Chr 2) marks the location of Apbb1ip. **(C)** Pld4 has the highest module membership within the salmon module. Pld4 is located on Chr 12 (marked with a green triangle), and its expression in the hippocampus maps as a trans-eQTL to the Apbb1ib locus.

### Age-related association dynamics of co-expressed gene networks

We next applied differential correlation analysis (DCA) to examine correlation dynamics between MEs in relation to ageing. Here, we considered the MEs as representative proxies for co-expressed transcript networks. Unlike the DEA and WGCNA, which tested for linear associations across the full age spectrum, this DCA assessed how correlations change between age categories. This provided a complementary perspective of how associated modules of gene expression become altered with age. First, the 33 MEs were tested for pairwise differential correlation between age categories: young and aged (**Table 1**), which resulted in 528 paired comparisons. After multiple testing correction, 45 ME pairs (8.5%) were significantly differentially correlated between the age groups (**Figure 4A & Data S5**). As described previously, the modules were annotated based on GO enrichment analyses of module members (**Supplementary Figures 5-7, Data S3 & S4**). Focussing on the salmon (immune response) ME, it was differentially correlated with three other MEs: white (waste disposal), black (protein folding) and sky blue (RNA splicing), between young and aged mice. For the white and black modules this corresponded to a gain in association, and for the sky blue module, a flip in the direction of association (**Figure 4A**). For example, the association between the salmon (immune response) and white (waste disposal) modules within young mice was minimal (*ρ* = 0.15) and this association strengthened in the direction of a negative correlation in aged mice (*ρ* =-0.35; **Figure 4B**), a significant change in correlation (*p* = 2.37 x 10^-3^; **Data S5**). There was no evidence of a correlation between these modules in mice within the mid aged category (*ρ* =-0.01; **Figure 4B**). This gain of association, and other examples illustrated in the network (**Figure 4A**), highlights the connectivity dynamics of functional components within hippocampal gene expression during the ageing process. Moreover, this analysis supports that systems-level coordination in biological processes are an important aspect of age-related gene expression signatures.

**Figure 4:**
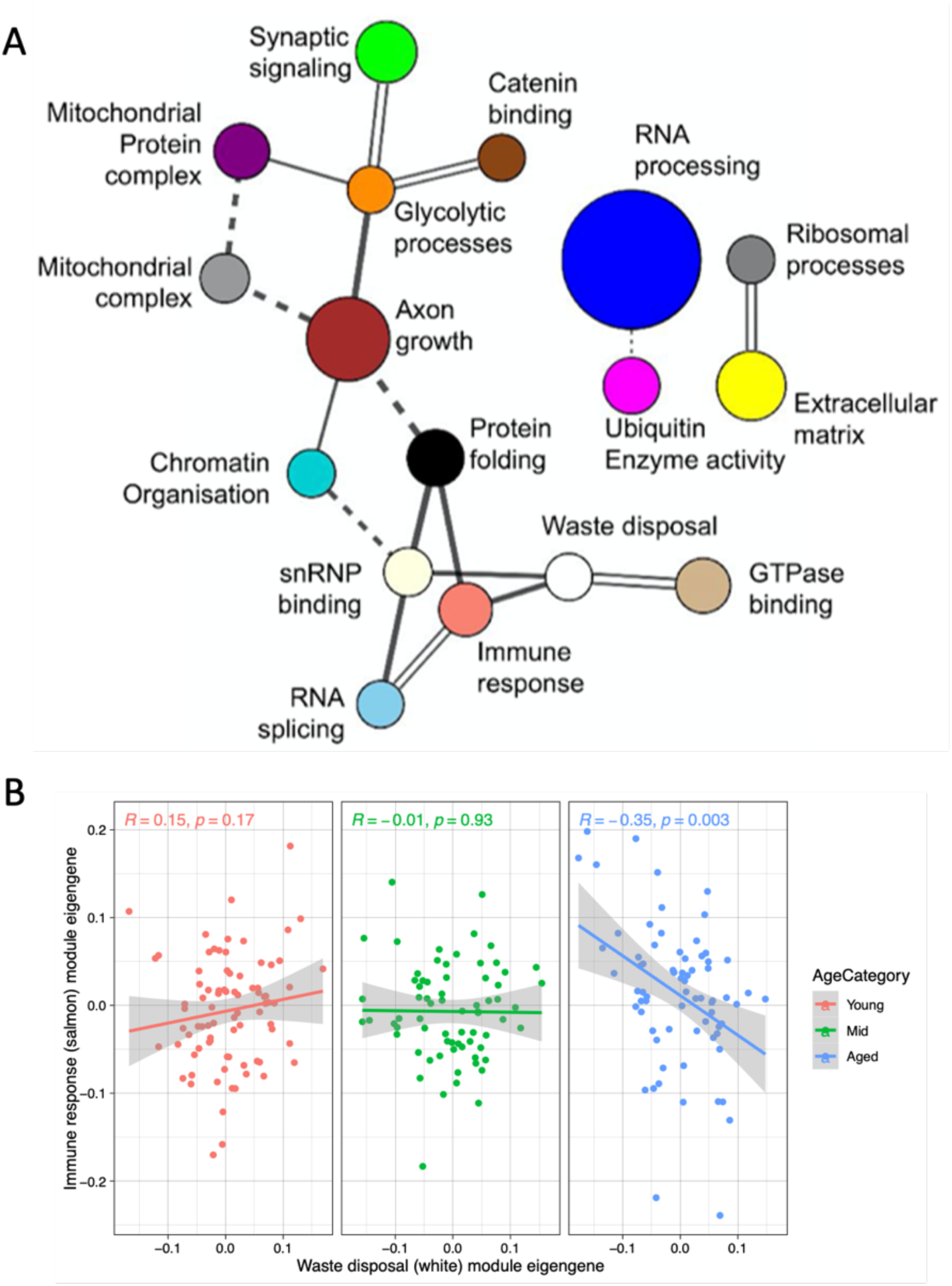
Differential correlation of co-expressed hippocampal gene modules across age. **(A)** Dynamic network of module eigengenes (MEs) annotated with enriched biological functions, derived from prior WGCNA. Edges represent differential correlation relationships between co-expressed modules in aged versus young mice. Solid lines represent a gain of association between corresponding nodes in aged mice. Dashed lines represent a loss of association and double lines represent a flip in the direction of the association, in aged mice. Node size corresponds to module size. **(B)** Scatter plot of the association between immune response (salmon) ME and waste disposal (white) ME for each sample in each age category. Relationships assessed using Spearman’s rank correlation.

### *Ywhab* highly differentially correlated between young and aged mice with implications for adapter:client protein relationships

To further investigate dynamic changes in co-expression during ageing, we performed DCA at the transcript-level. For the 26,273 transcripts, this resulted in ∼345 million pairs of gene correlations within each age group. At an FDR of 5%, 4515 pairs of transcripts were differentially correlated between young and aged mice (**Data S6**). These differential correlations represented 3056 genes, most only differentially correlated with one other gene (n = 1850; **Figure 5A**). Exceptions included *Ywhab*, *Rpl17, Ube2d1, Gabrr2* and *Spryd4* which were the top differentially correlated genes between young and aged mice. The expression of *Ywhab* was differentially correlated with 74 other transcripts, accounting for 71 different genes (**Figure 5A**), hinting at a potential regulatory dysfunction of *Ywhab* with ageing. For instance, in young animals, *Ywhab* was strongly positively correlated with *Map2k4* (*ρ* = 0.63), which changed to an inverse correlation in aged animals (*ρ* =-0.34), a statistically significant loss in association (*p* = 1.84 x 10^-10^; **Figure 5B & C**). A similar loss of correlation was observed for 33 of the 71 differentially correlated genes, largely from a strong positive correlation in young animals to a weak inverse correlation in aged animals. A gain in correlation was observed for 13 genes (e.g. weak or no correlation in young animals to a stronger correlation in aged animals, such as for *Acat1*), and a flip in correlation was evident for 25 genes, for example for *Ptma* (**Figure 5B & C**).

**Figure 5:**
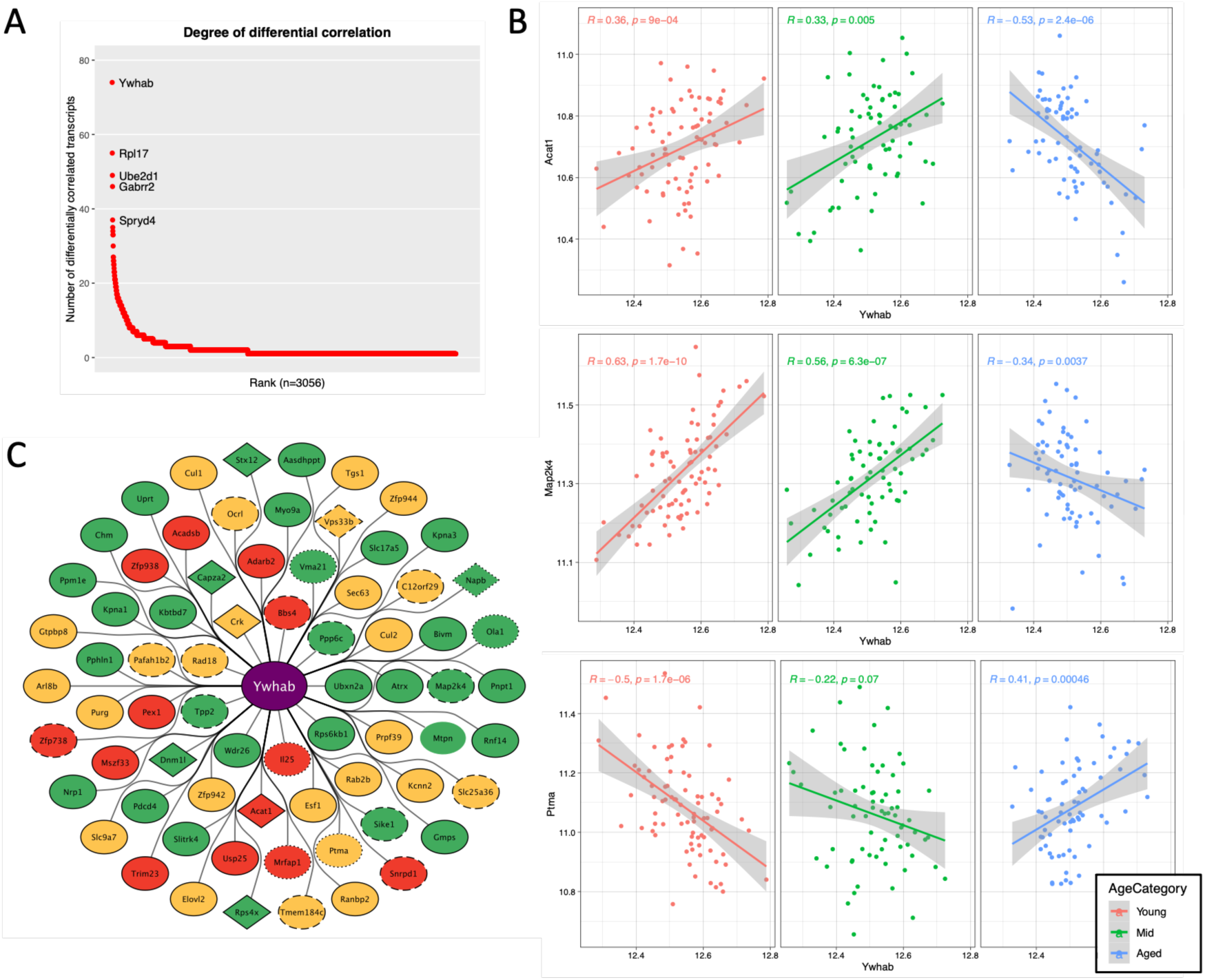
Ywhab is highly differentially correlated in the hippocampus during ageing. **(A)** Rank plot illustrating the degree of differential correlation within the hippocampal transcriptome between young and aged mice. Greater than 3000 genes are differentially correlated and these are ordered based on their number of differentially correlated genes. The top 5 differentially correlated genes are labelled. **(B)** Scatterplots and Spearman correlations of gene expression of 3 genes differentially correlated with Ywhab: Acat1, Map2k4 and Ptma, across the 3 age categories: Young, Mid and Aged. **(C)** Network graph summarising the extent of genes differentially correlated with Ywhab between young and aged mice. Node colour represents the change in association: gain of association (red), loss of association (green), flip in the association direction (yellow). The node border corresponds to whether the protein product of Ywhab (a 14-3-3 protein) is predicted to bind the protein product of the node transcript, assessed using 3 algorithms within the 14-3-3-Pred tool^41^: not predicted to bind (no border), predicted to bind by 1 model (dotted border), predicted to bind by 2 models (dashed border), predicted to bind by all 3 models (filled border). The node shape represents whether there is protein interaction evidence supporting the relationship: no protein interaction evidence (oval), protein interaction evidence (diamond). Transcript/exon level data was collapsed to gene level information.

*Ywhab* encodes the 14-3-3-beta/alpha protein, a highly conserved adapter molecule which binds phosphosites of client proteins to regulate many aspects of their behaviour, such as enzymatic activity, protein folding and protein interaction potential^46^. With this in mind, utilising the 14-3-3-Pred tool^41^, we explored whether the protein products of the differentially correlated transcripts of *Ywhab* are likely to be client proteins, based on multiple sequence-based machine learning (ML) prediction models. Interestingly, 70% of the proteins encoded by genes differentially correlated contain likely 14-3-3 binding motifs as predicted by three distinct ML models (**Figure 5C & Data S7**). Only one protein, myotrophin (Mtpn), did not contain a likely 14-3-3 binding sequence, across either of the three models. To further assess whether these proteins are indeed 14-3-3 clients, reported protein-protein interaction data was exploited using MIST^42^. Eight of these proteins (Acat1, Capza2, Crk, Dnm1l, Napb, Rps4x, Stx12, Vps33b; depicted with diamond nodes in **Figure 5C**) are reported to interact with 14-3-3-beta/alpha. Upon assessing GO Cellular Component annotations assigned to these 8 proteins (**Data S7**), dysregulation of these adapter:client interactions could have broad implications for subcellular homeostasis, in particular relating to membrane biology. Altogether, these results showcase the dynamic nature of relationships in hippocampal gene expression across age, both at a co-expression cluster and individual gene level.

## Discussion

In this work we leveraged a hippocampal gene expression dataset generated from the BXD mouse reference population^20^. This inbred population combines the benefits of recombination (large genetic diversity, yet high mapping power due to only two progenitor strains) and homozygosity (owing to the inbred status of each strain). This allows for comparison of phenotypes measured in the same strains without needing to be from the same animals, or even the same study—thus enabling powerful meta-analyses as presented in this work. Our goal was to apply complementary strategies to explore age-dependent dynamics in the hippocampal transcriptome, with an emphasis on maximising dataset utility. Through a systems genetics approach^28^, we interrogated these data with differential expression, differential correlation, co-expression network and quantitative trait loci analyses, further supplemented by enrichment and focussed data mining approaches.

First, we corroborated that immune responses and microglial involvement are core features of hippocampal ageing, in support of existing literature^44,47,48^. Since our study was performed in a genetically diverse but normative cohort of aged mice rather than an age-related disease model, the increase in immune signatures with ageing indicates that neuroinflammation is a prominent and representative feature of the ageing brain^49^. This immune profile likely contributes to the increased susceptibility to neurodegenerative diseases^50,2^ and underscores the need for a better understanding of the molecular underpinnings of immune responses in the ageing brain.

Importantly, we identified specific microglial subtypes or cell states that are major cellular drivers of the gene expression landscape observed in the aged hippocampus. This is potentially due to an increase in abundance of these subtypes/states, for example the recently described age-dependent microglia^44^. However, due to microglial heterogeneity^51^, it is likely that other glial subtypes (and other brain cell types not accounted for in our analysis) contribute to brain ageing^52^. In future work, more comprehensive single-cell datasets may reveal additional age-associated cell type enrichments and help address whether cell subtype compositions change during ageing. This provides an example of how the utility of archived datasets can be enhanced by new, more technologically advanced data and methods.

Since transcripts and their protein products function within an interconnected coregulated cellular landscape, we assessed co-expression of transcripts in the ageing hippocampus and the age-related dynamics of co-expressed modules. These analyses highlighted significant involvement of immune response and myelination co-expression gene signatures during ageing. Further analysis indicated that this immune response signal is likely derived from microglia and perivascular macrophages within the tissue, reinforcing the role of microglial immune responses in hippocampal ageing. Moreover, our findings support perivascular macrophages as important players in this process^53^. Interestingly, the immune response module was differentially correlated with three other co-expression modules: waste disposal, protein folding and RNA splicing, which are processes previously implicated in ageing^54,55^. However, this result suggests there are changes in how these processes are coordinated (directly or indirectly) during ageing and enables module relationships to be assessed at specific ageing timepoints. As an example, the gain in association between immune response and waste disposal modules from a slight positive correlation to a significant inverse correlation at the aged timepoint, implies gene expression programs underlying waste disposal processes become less active as immune response gene expression increases. This shift in correlation suggests an age-driven collapse of immune–proteostasis coupling that may contribute to deficits in microglial clearance of protein aggregates observed in neurodegenerative diseases^56^.

Whether there are mechanistic links underlying these dynamics is an intriguing route for further investigation.

Moreover, genetic regulation of the immune response module mapped to the *Apbb1ip* locus on Chr 2. *Apbb1ip* is a member of the immune response (salmon) module and encodes the amyloid beta precursor protein binding family B member 1 interacting protein. This protein is linked to T cell activation in the periphery^57^ and is emerging as a core component of microglial expression signatures^58,59,60^. Furthermore, genetic association studies in humans implicate *APBB1IP* in Alzheimer’s disease and schizophrenia^61,62^. Our study provides evidence that genetic variants that have a *cis*-regulatory effect on *Apbb1ip* expression also exert a *trans-*regulatory effect on co-expressed genes within the age-related immune response module. Specifically, the expression of immunoregulatory exonuclease *Pld4*^63^, which exhibits the highest module membership within the immune response (salmon) module, mapped as a *trans-*eQTL to the *Apbb1ip* locus, suggesting a *trans*-modulatory influence by *Apbb1ip*.

Further resolution on differential correlations between young and aged mice revealed *Ywhab* as a top hit among >3000 other differentially correlated genes, highlighting the dynamic nature of gene expression in the ageing hippocampus. *Ywhab* encodes 14-3-3-beta/alpha, a highly conserved adaptor protein and one of seven 14-3-3 protein isoforms important for regulating phosphosites of client proteins. 14-3-3 proteins influence a diverse range of signalling events^64^ including modulation of inflammation^65^. Of note, 14-3-3s are abundantly and ubiquitously expressed in the brain and have been implicated in many neurodegenerative diseases^46,66,67^. These chaperone proteins bind recognition motifs of clients and through protein interaction analysis we predicted that the majority of proteins encoded by the differentially correlated transcripts are likely binding partners, with reported protein interaction evidence for eight of them. This provided intriguing evidence at the protein level for the functional consequences of *Ywhab* differential correlation. Due to the high degree of differential correlation *Ywhab* exhibits during ageing and its vast signalling network potential, 14-3-3-beta/alpha is an important hub which may underlie widespread age-associated molecular phenotypes and/or dysfunction in the hippocampus, including processes relating to immune function.

Collectively, our research provides insight into the complex patterns of hippocampal gene expression changes during ageing. We demonstrated a strong involvement of immune responses in the ageing hippocampus and captured gene expression dynamics at transcript and functional module resolution, providing novel systems-level perspectives on the interplay of gene expression changes at various ageing timepoints. Our study implicates *Apbb1ip* as a key genetic modulator of immune responses and microglial activity during normative ageing, which warrants further investigation.

## Acknowledgements

We would like to thank the Summer School *Systems Genetics of Neural Ageing* (August 2022) for bringing the authors together and triggering this collaboration. We would like to acknowledge the funding for this summer school from the e:Med Systems Medicine programme of the BMBF (*Bundesministerium für Bildung und Forschung*; German Ministry of Education and Research) to RWO. This research was funded in part by Aligning Science Across Parkinson’s [ASAP-000509] through the Michael J. Fox Foundation for Parkinson’s Research (MJFF), by supporting JET.

## Supplementary Figures

**Supplementary Figure 1.**
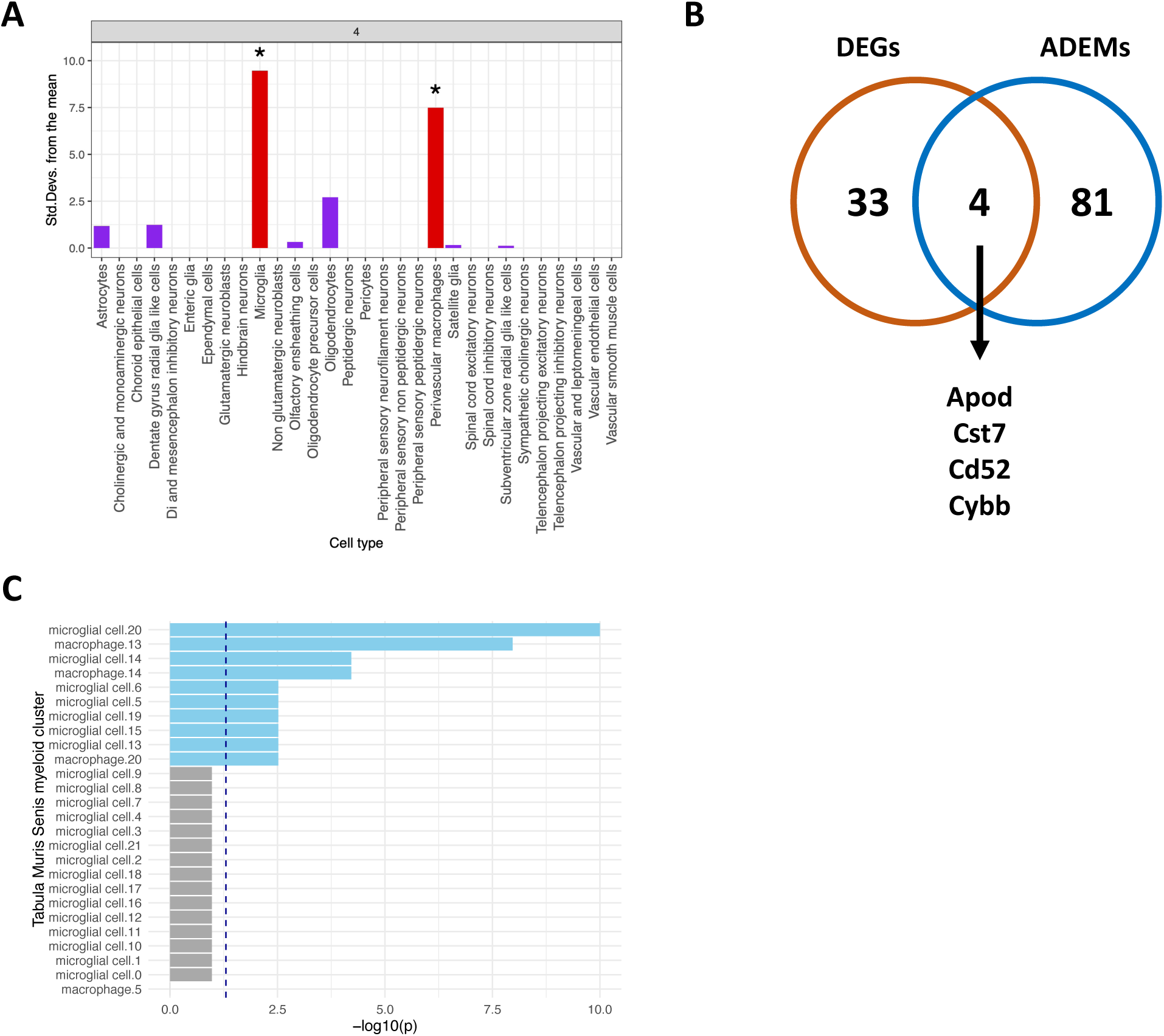
Age-associated transcriptomic changes in the hippocampus are enriched within microglia and perivascular macrophages. **(A)** Barplot showing cell type enrichments calculated using EWCE^30^. EWCE was performed using DEGs from the ageing mouse hippocampus and a mouse whole nervous system single-cell dataset^43^. Microglia and perivascular macrophages were enriched for the DEGs after FDR correction. **(B)** Intersection of DEGs with age-associated microglia (ADEM) marker gene list^44^. **(C)** ADEM marker genes are overrepresented in microglia cluster 20, and to a lesser extent several other microglia and macrophage clusters, as defined within the Tabula Muris Senis study^31^.

**Supplementary Figure 2.**
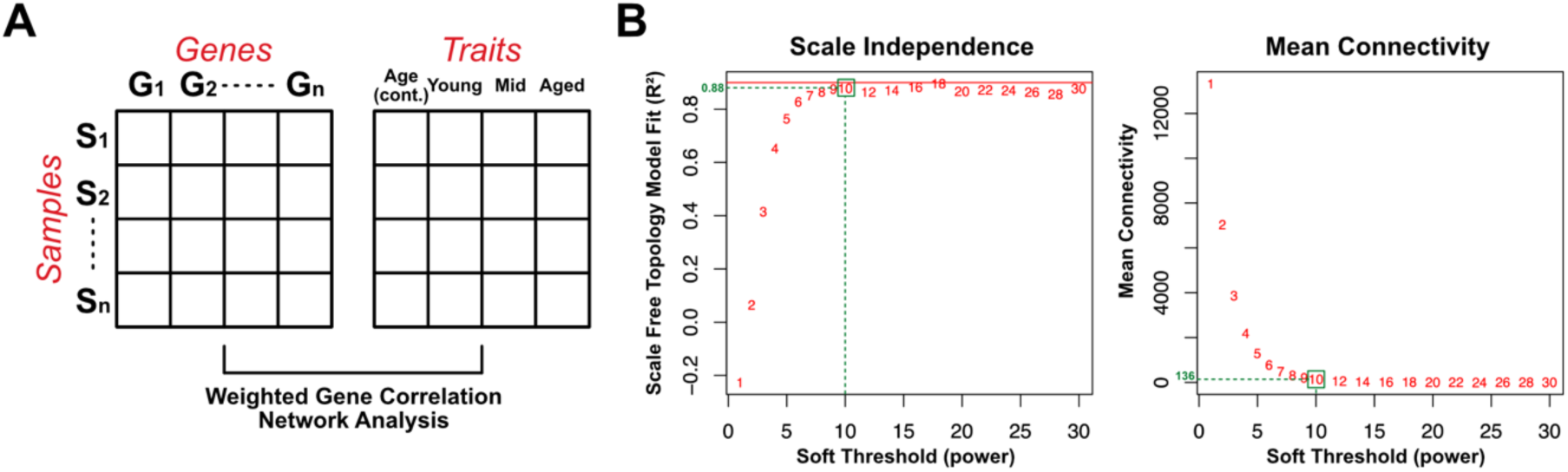
Weighted gene co-expression analysis of aged hippocampal samples from BXD mice. **(A)** Overview of expression data organised as a gene by sample matrix of 26,273 transcripts across 223 samples. The external traits used were age (as a vector of discrete data points for each sample) and age binned into three groups with binary values – Young (< 480 days), Mid (480 - 600 days) and Aged (> 600 days). **(B)** Plots of scale-free topology fit index and mean connectivity as a function of soft thresholding power.

**Supplementary Figure 3.**
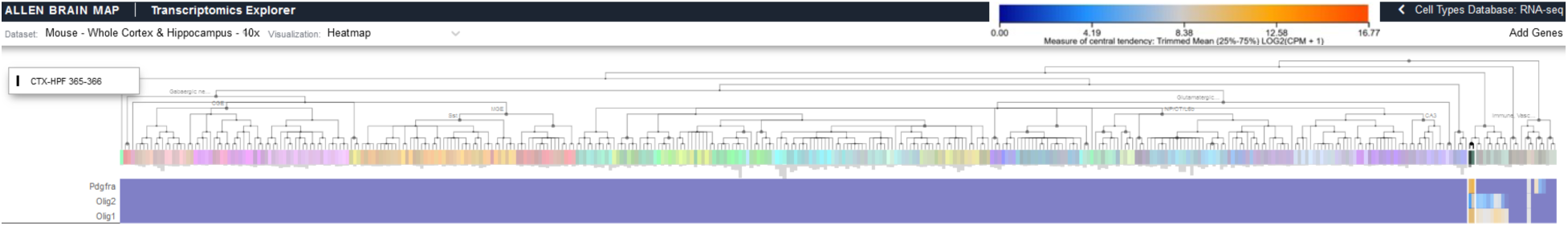
Expression of OPC marker genes. Gene expression of Pdgfra, Olig2 and Olig1 in cluster 365 of the Mouse Whole Cortex & Hippocampus 10x dataset from the Allen Brain Map^34^.

**Supplementary Figure 4.**
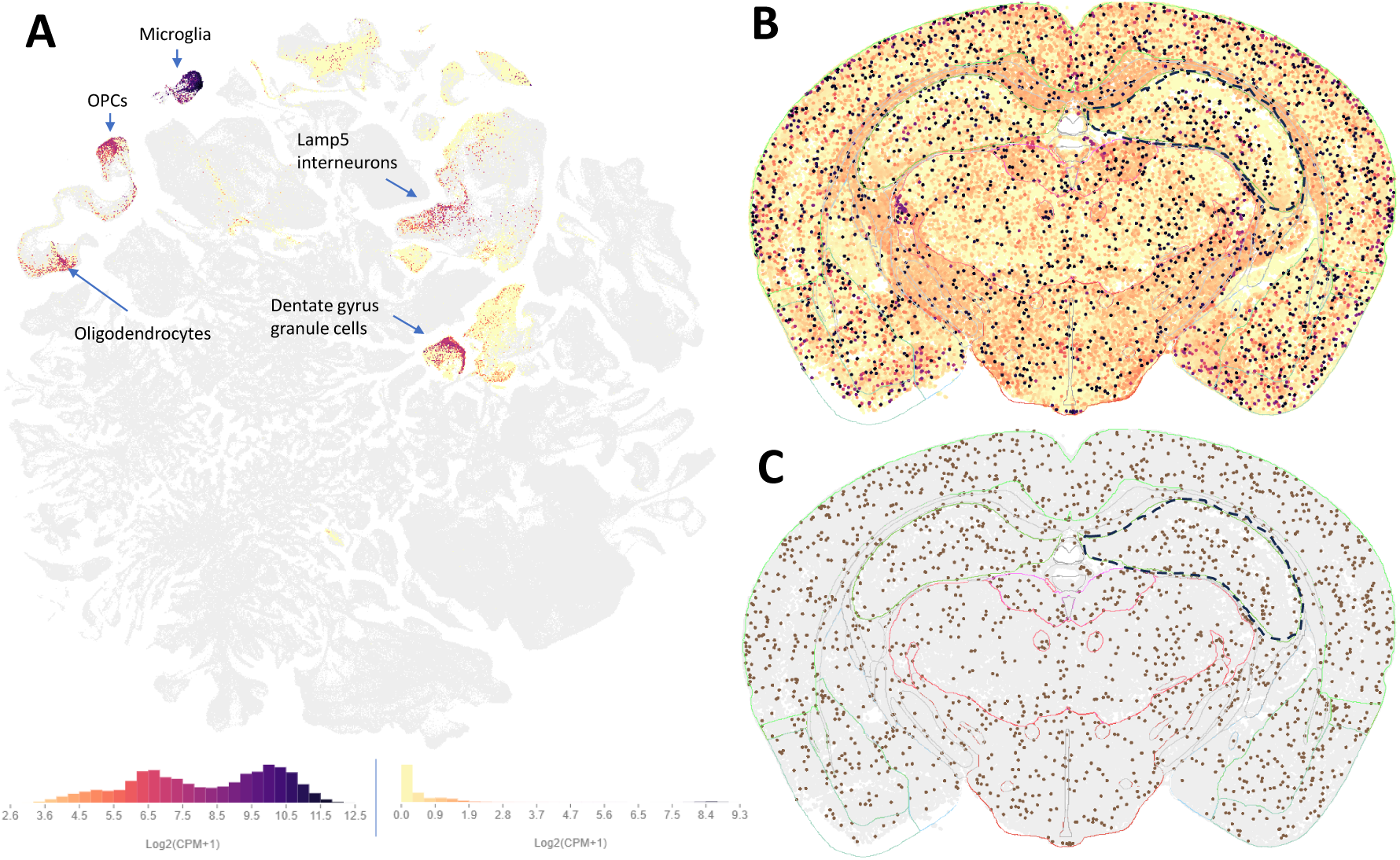
**Apbb1ip most highly expressed in microglia in the mouse hippocampus**. **(A)** Expression levels of Apbb1ip in snRNA-seq data from the mouse hippocampus, overlaid on a UMAP of the whole mouse brain. Left colour bar below shows expression levels as a density plot. Expression is by far the highest in microglia and shows low to moderate expression in a subset of other labelled cell types. **(B)** Expression levels of Apbb1ip in a representative tissue section (C57BL6J-638850.36) based on spatial transcriptomics (MERFISH). A small number of cells show very high expression while the remaining cells show no to very low expression across the brain, including in hippocampus (surrounded by black dashed line). **(C)** Colour coding of same tissue section to highlight microglia (brown; all other cells in grey) show overlap with cells showing high expression of Apbb1ip, strengthening results from scRNA-seq. All images shown from the Allen Brain Cell Atlas (RRID:SCR_024440); https://portal.brain-map.org/atlases-and-data/bkp/abc-atlas. All data from^68^.

**Supplementary Figure 5.**
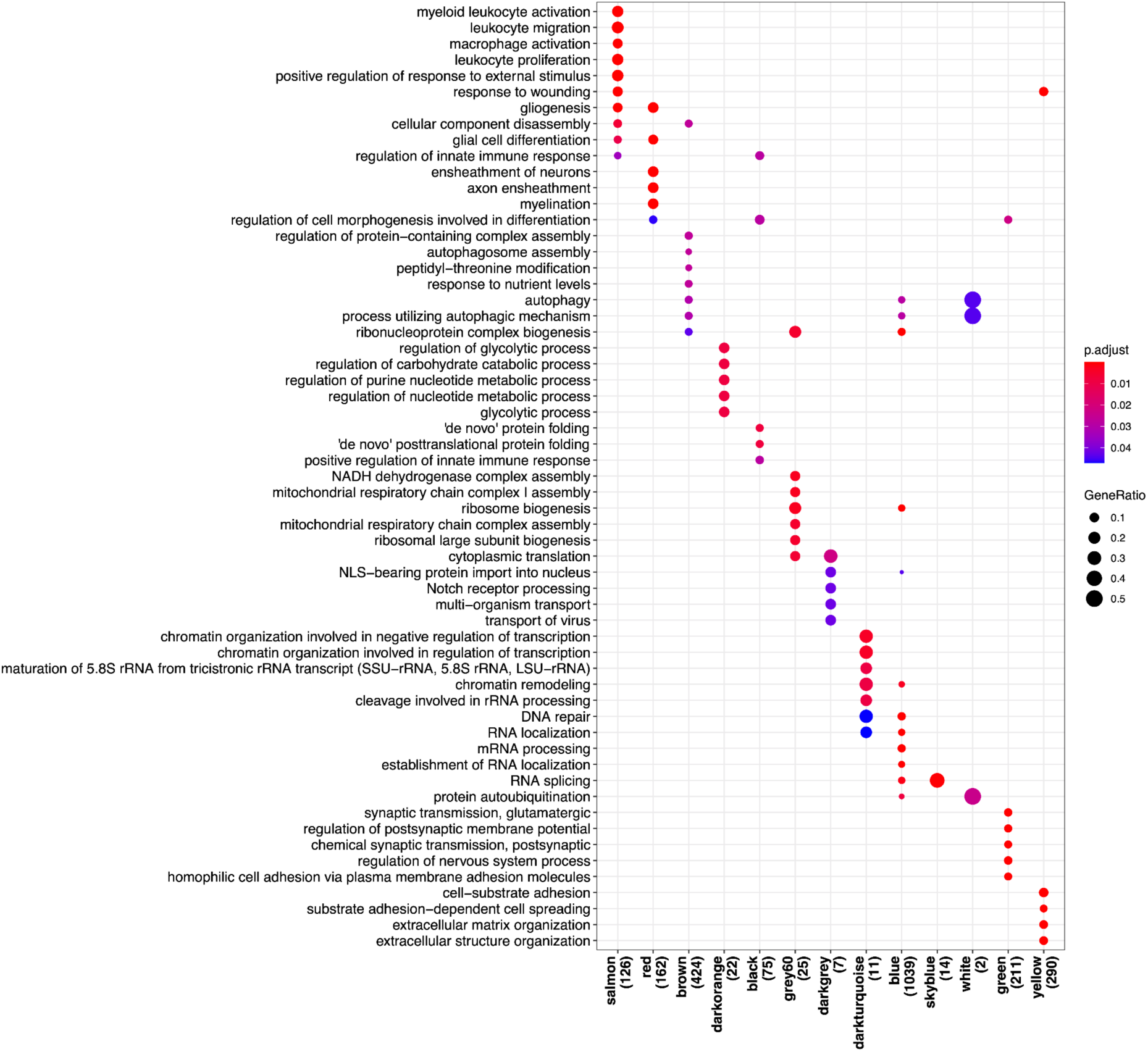
Gene ontology (biological process) enrichment analysis for selected WGCNA modules. Gene ontology annotations listed on the y-axis and module annotations (with number of annotated genes in module) on the x-axis. Datapoint colour represents adjusted p-value of enrichment result and size represents gene ratio, i.e., the proportion of queried genes contributing to the enrichment result.

**Supplementary Figure 6.**
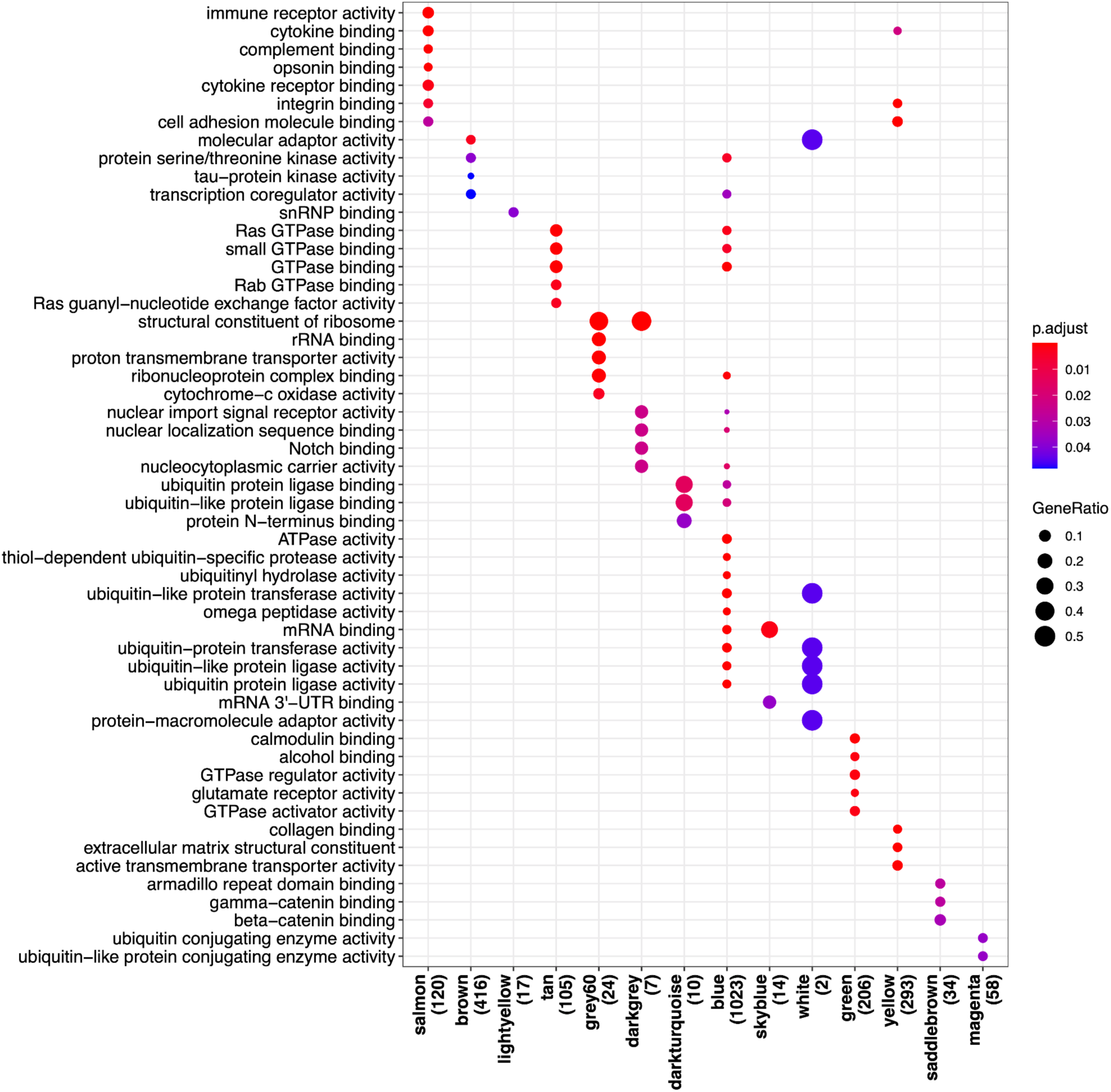
Gene ontology (molecular function) enrichment analysis for selected WGCNA modules. Gene ontology annotations listed on the y-axis and module annotations (with number of annotated genes in module) on the x-axis. Datapoint colour represents adjusted p-value of enrichment result and size represents gene ratio, i.e., the proportion of queried genes contributing to the enrichment result.

**Supplementary Figure 7.**
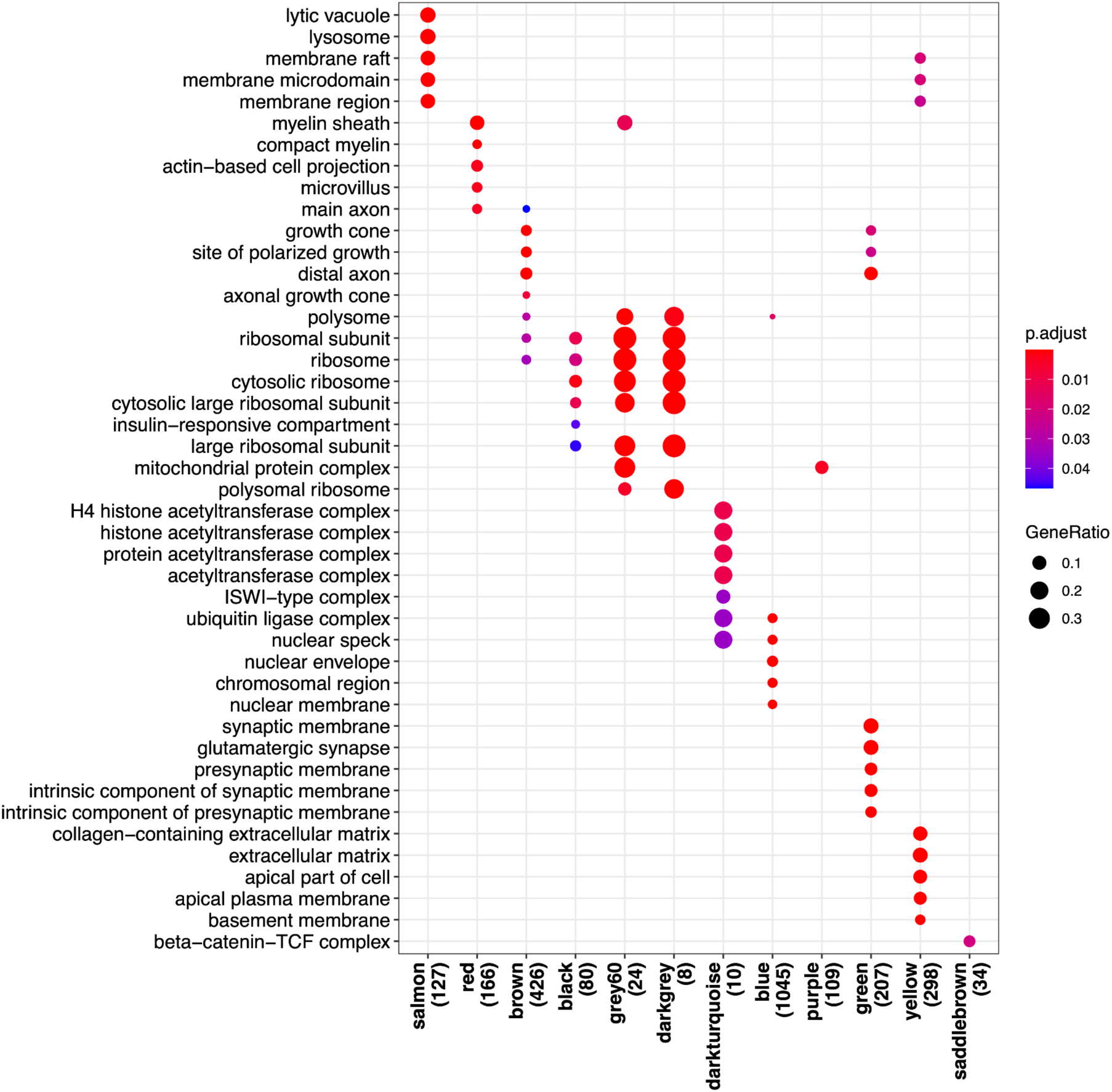
Gene ontology (cellular component) enrichment analysis for selected WGCNA modules. Gene ontology annotations listed on the y-axis and module annotations (with number of annotated genes in module) on the x-axis. Datapoint colour represents adjusted p-value of enrichment result and size represents gene ratio, i.e., the proportion of queried genes contributing to the enrichment result.

## Supplementary Files

Data S1 – Sample demographics and sample-level WGCNA principal component loadings module eigengene values

Data S2 – Differential expression and WGCNA results Data S3 – Description of WGCNA modules

Data S4 – Gene ontology functional enrichment analysis results for WGCNA modules Data S5 – Module eigengene differential correlation analysis results

Data S6 – Transcript-level differential correlation analysis significant results

Data S7 – Ywhab differentially correlated genes, 14-3-3 binding prediction and protein interaction evidence

