## Supplemental Appendix 1 for "Genetic modulation of immune gene co-expression in the aged mouse hippocampus by the *Apbb1ip* locus": differential_correlation_on_eigen_clean.R

```
# Load necessary libraries
library(igraph)
library(ggplot2)
library(openxlsx)

# Function to normalize values between 0 and 1
range01 <- function(x) {
  (x - min(x)) / (max(x) - min(x))
}

# Load module eigengene vectors from CSV file
eigen <- read.csv("data/moduleEigengeneVectors_allModules.csv", stringsAsFactors = FALSE)
rownames(eigen) <- eigen$X  # Set row names to the values in the 'X' column
eigen$X <- NULL             # Remove the 'X' column
eigen <- t(eigen)           # Transpose the matrix

# Check dimensions of the eigen data
dim(eigen)

# Load sample annotations
samplesheet <- read.csv("data/SampleAnnotations_AgedHippo_withAgeBins.csv", stringsAsFactors = FALSE)

# Match samplesheet to the columns in eigen data
samplesheet <- samplesheet[match(colnames(eigen), samplesheet$Strain), ]

# Clean memory
gc()

# Prepare data for analysis
eigen <- eigen[, samplesheet$Strain]        # Reorder columns in eigen to match samplesheet
samplesheet$colnames <- colnames(eigen)     # Add column names to samplesheet

# Run differential correlation analysis using module eigengenes
difcor_eigen <- difcorRun(
  data = eigen,
  samplesheet = samplesheet,
  contrasCol = "Age_3bins",
  contrast = c("Early", "Late"),
  method = "spearman",
  padjust = "perm",
  Nperm = 1000
)

# Adjust p-values using Benjamini-Hochberg method
difcor_eigen$fdr <- p.adjust(difcor_eigen$P, method = "BH")

# View the first few rows of the results
head(difcor_eigen)

# Filter results where GeneB is 'MEsalmon'
difcor_eigen[difcor_eigen$GeneB == "MEsalmon", ]

# Get the number of significant results with permuted p-value <= 0.05
significant_results_dim <- dim(difcor_eigen[difcor_eigen$perm_P <= 0.05, ])
print(significant_results_dim)

# Calculate the proportion of significant results
significant_proportion <- 45 / 528
print(significant_proportion)

# Write the differential correlation results to an Excel file
write.xlsx(difcor_eigen, file = "Module_DCA_results.xlsx")

# Filter significantly different correlations with permuted p-value <= 0.01
superSign <- difcor_eigen[difcor_eigen$perm_P <= 0.01, ]

# Calculate the difference in correlation between conditions
superSign$delta <- superSign$AD_cor - superSign$ct_cor

# Label the type of change based on delta
superSign$type <- ifelse(superSign$delta > 0, "Increase", "Decrease")

# Check the dimensions of the significant results
dim(superSign)

# View entries where GeneA or GeneB is 'MEsalmon'
superSign[superSign$GeneA == "MEsalmon" | superSign$GeneB == "MEsalmon", ]

# Classify changes as 'Gain', 'Loss', or 'Flip' based on correlation differences
superSign$type <- ifelse(
  abs(superSign$ct_cor) - abs(superSign$AD_cor) < 0, "Gain", "Loss"
)
superSign$type <- ifelse(
  abs(abs(superSign$ct_cor) - abs(superSign$AD_cor)) <= 0.10, "Flip", superSign$type
)

# Define line types and colors for plotting based on the type of change
superSign$linetype <- unname(c("Gain" = "solid", "Loss" = "dotted", "Flip" = "solid")[superSign$type])
superSign$linecolor <- unname(c("Gain" = "black", "Loss" = "black", "Flip" = "red")[superSign$type])

# Calculate edge weights for network plotting
superSign$weight <- range01(abs(superSign$delta)) * 5

# Write the significant results to a CSV file for network visualization
write.csv(superSign, file = "Module_graph.csv", quote = FALSE, row.names = FALSE)

# Write the list of unique modules involved in significant changes
unique_modules <- unique(c(superSign$GeneA, superSign$GeneB))
write.csv(unique_modules, file = "DCA_modules.csv", quote = FALSE, row.names = FALSE)

# Create a network from the significant results
network <- makeNetwork(superSign)

# Calculate centrality measures in the network
centrality <- getCentrality(network)

# View the top 100 nodes by degree centrality
head(centrality, 100)

# Prepare data for network plotting
early <- superSign[, c("GeneA", "GeneB", "delta", "type")]

# Create a data frame of links (edges) for the network
links <- data.frame(
  source = as.character(early$GeneA),
  target = as.character(early$GeneB),
  weight = superSign$delta,
  type = superSign$type
)

# Create a data frame of nodes (vertices) for the network
genes <- unique(c(links$source, links$target))
nodes <- data.frame(nodes = genes)

# Create the network graph using igraph
network <- graph_from_data_frame(d = links, vertices = nodes, directed = FALSE)

# Plot the network using igraph
plot(
  network,
  layout = layout_with_dh(network),
  edge.color = superSign$linecolor,
  edge.lty = superSign$linetype,
  edge.width = range01(abs(links$weight)) * 5,
  vertex.label.cex = 0.7,
  vertex.size = 5
)

# View the last few rows of significant results
tail(superSign)

# Load gene module membership data
membership <- read.csv("data/geneModuleMembership_allModules.csv", stringsAsFactors = FALSE)
rownames(membership) <- membership$X  # Set row names
membership$X <- NULL                  # Remove 'X' column
membership <- t(membership)           # Transpose the matrix

# View the first 10 columns of membership data
membership[, 1:10]

# Extract genes with high membership (>= 0.6) in 'brown' and 'magenta' modules
MEBrown_genes <- colnames(membership)[membership["MMbrown", ] >= 0.6]
MEmagenta_genes <- colnames(membership)[membership["MMmagenta", ] >= 0.6]

# Clean gene names by removing any suffix after a period
MEBrown_genes <- sapply(strsplit(MEBrown_genes, split = "[.]"), function(x) x[1])
MEmagenta_genes <- sapply(strsplit(MEmagenta_genes, split = "[.]"), function(x) x[1])

# Write the gene lists to CSV files
write.csv2(MEBrown_genes, file = "MEBrown_genes.csv", quote = FALSE, row.names = FALSE)
write.csv2(MEmagenta_genes, file = "MEmagenta_genes.csv", quote = FALSE, row.names = FALSE)

# Define module eigengenes for plotting
geneA <- "MEtan"
geneB <- "MEwhite"

# Prepare data for plotting module eigengenes across age groups
plotData <- data.frame(
  geneA = eigen[geneA, ],
  geneB = eigen[geneB, ],
  ageGroups = factor(samplesheet$Age_3bins, levels = c("Early", "Middle", "Late"))
)

# Calculate Spearman correlation between geneA and geneB in the 'Late' age group
correlation_late <- cor(
  plotData$geneA[plotData$ageGroups == "Late"],
  plotData$geneB[plotData$ageGroups == "Late"],
  method = "spearman"
)
print(paste("Spearman correlation in Late group:", correlation_late))

# Plot the relationship between geneA and geneB using ggplot2
ggplot(plotData, aes(x = geneA, y = geneB, color = ageGroups)) +
  geom_point() +
  facet_grid(~ ageGroups) +
  geom_smooth(method = "lm", se = FALSE) +
  labs(
    x = geneA,
    y = geneB,
    title = paste("Expression of", geneA, "vs", geneB, "by Age Group")
  ) +
  theme_minimal() +
  xlim(-0.2, 0.15)

# Note: Variables 'brown_darkorange' and 'brown_black' are not defined in the provided code.
# If you have these plots defined elsewhere, you can combine them using patchwork or gridExtra.

# Reload samplesheet if needed
samplesheet <- read.csv("data/SampleAnnotations_AgedHippo_withAgeBins.csv", stringsAsFactors = FALSE)

# Check dimensions of samplesheet
dim(samplesheet)

# Display the distribution of sexes
table(samplesheet$Sex)

# Summary statistics of the 'Age' variable
summary(samplesheet$Age)

# Summary statistics of 'Age' for male samples
summary(samplesheet[samplesheet$Sex == "M", "Age"])
```
