## Supplemental Appendix 1 for "Genetic modulation of immune gene co-expression in the aged mouse hippocampus by the *Apbb1ip* locus": Differential_expression.R

```
library(rio)

data <- readRDS("GroupC_BXDAgedHippocampus_AffyExpression.rds")

expr <- data$Expression
geneInfo <- data$geneInfo

samplesheet <- import("SampleAnnotations_AgedHippo_withAgeBins.csv")
samplesheet <- samplesheet[match(colnames(expr),samplesheet$Strain),]

allRes <- lapply(1:nrow(expr),function(i){
  message(i)
  tmpData <- data.frame(gene = expr[i,],
                        age = samplesheet$Age,
                        sex = samplesheet$Sex,
                        batch = samplesheet$Batch)
  
  res <- lm(formula = "gene ~ age + sex + batch", data = tmpData)
  out <- summary(res)
  out <- out$coefficients[2,]
  return(out)
}) |> do.call(what = "rbind")

rownames(allRes) <- rownames(expr)
allRes <- as.data.frame(allRes)
allRes$fdr <- p.adjust(allRes$`Pr(>|t|)`,method = "BH")
allRes <- allRes[order(allRes$fdr),]

library(ggplot2)
ggplot(allRes, aes(Estimate, -log10(`Pr(>|t|)`))) + geom_point() + theme_minimal()

table(allRes$fdr<=0.05)
```
