## Supplemental Appendix 1 for "Genetic modulation of immune gene co-expression in the aged mouse hippocampus by the *Apbb1ip* locus": preprocessData_WGCNA.R

```
## Libraries
{
  pkg <- c("foreach",
           "GenomicRanges",
           "GenomicFeatures",
           "gplots", 
           "ggplot2",
           "data.table",
           "sva",
           "doParallel",
           "BiocParallel",
           "DESeq2",
           "RUVSeq",
           "ggpubr",
           "cowplot",
           "GGally",
           "caret", 
           "vsn",
           "IHW",
           "pheatmap",
           "RColorBrewer",
           "tibble",
           "EnhancedVolcano",
           "variancePartition",
           "PCAtools",
           "VennDiagram")
  lapply(pkg, function(x) {
    if(!require(x,character.only = TRUE)) install.packages(x,character.only = TRUE)
  })
}       

####################################################################################################
## Loading and curating data
####################################################################################################

resultsPath <- "/Users/methiaa/Downloads/GroupC_BulkRNA_WGCNA/"

## Loading expression matrix
norm_data <- readRDS(paste(resultsPath, "cts/GroupC_BXDAgedHippocampus_AffyExpression.rds", sep=""))
norm_matrix <- as.data.frame(norm_data$Expression)

## Loading covariate data
meta_data <- read.table(file=paste(resultsPath, "normalization/SampleAnnotations_AgedHippo_withAgeBins.csv", sep=""),
                        header=TRUE,
                        sep=",",
                        stringsAsFactors=TRUE)
row.names(meta_data) <- meta_data$ID
meta_data <- meta_data[,-c(1:3)]
meta_data$Sex <- factor(meta_data$Sex)
meta_data$Batch <- factor(meta_data$Batch)
meta_data$Strain <- factor(meta_data$Strain)
#meta_data$Age <- factor(meta_data$Age)
meta_data$Age_3bins <- factor(meta_data$Age_3bins)

meta_data.0 <- meta_data

norm_matrix <- norm_matrix[,match(rownames(meta_data), colnames(norm_matrix))]

### Variance partition

# Specify variables to consider
# (Age is continuous so model it as a fixed effect
# Batch and Sex are both categorical, so model them as random effects)
form <- ~ Age + (1|Sex) + (1|Batch)

# Fit model and extract results
varPart <- fitExtractVarPartModel(norm_matrix, form, meta_data)

# sort variables (i.e. columns) by median fraction of variance explained
vp <- sortCols(varPart)
# # Bar plot of variance fractions for the first 10 genes
# plotPercentBars(vp[1:10,])
# violin plot of contribution of each variable to total variance
plotVarPart(vp)

### PCA plots
rownames(meta_data) == colnames(norm_matrix)
p <- PCAtools::pca(norm_matrix, metadata = meta_data)

pdf(file = paste(resultsPath,"normalization/PCA_before_removingBE.pdf",sep=""), height = 8, width = 10)
PCAtools::biplot(p, colby = 'Age_3bins', shape = "Sex", lab = "")
dev.off()

# ### Remove covariates
# 
# mod.matrix <- model.matrix(~Age_3bins, meta_data)
# 
# norm_matrix_covRem <- limma::removeBatchEffect(norm_matrix, covariates = meta_data[,c(1,3)], design = mod.matrix)
# plotPCA(norm_matrix_covRem, intgroup = "Age_3bins", ntop = 500) + 
#   geom_text(aes(label=colnames(count.data.0)),vjust=2) + 
#   labs(color = 'Age_3bins') + 
#   theme_pubr(legend = "right")

####################################################################################################

# Save corrected and normalized reads as csv file
cat("\t7) Saving normalized gene counts to csv file ...\n")
write.csv(norm_matrix, file=paste(resultsPath,"/normalization/normalized_counts.csv", sep = ''), row.names = T, quote = F, col.names = NA)
saveRDS(norm_matrix, file=paste(resultsPath,"/normalization/normalized_counts.rds", sep = ''))
```
