## Supplemental Appendix 1 for "Genetic modulation of immune gene co-expression in the aged mouse hippocampus by the *Apbb1ip* locus": run_differential_correlation_clean.R

```
# Load utility functions from an external script
source("utils.R")

# Load expression data from an RDS file
exprData <- readRDS("data/GroupC_BXDAgedHippocampus_AffyExpression.rds")
expr <- exprData$Expression   # Extract the expression matrix
geneInfo <- exprData$geneInfo # Extract gene information
rm(exprData)                  # Remove the loaded object to free memory

# Load sample annotations from a CSV file
samplesheet <- read.csv("data/SampleAnnotations_AgedHippo_withAgeBins.csv", stringsAsFactors = FALSE)

# Ensure that the samples in samplesheet match the columns in the expression data
samplesheet <- samplesheet[match(colnames(expr), samplesheet$Strain), ]

# Clean up memory
gc()

# Prepare expression data and samplesheet for analysis
expr <- expr[, samplesheet$Strain]           # Reorder columns in expr to match samplesheet
colnames(expr) <- make.names(colnames(expr)) # Make column names syntactically valid
samplesheet$colnames <- colnames(expr)       # Add column names to samplesheet

# Run differential correlation analysis using the first 10,000 genes
difcor_res <- difcorRun(
  data = expr[1:10000, ],
  samplesheet = samplesheet,
  contrasCol = "Age_3bins",
  contrast = c("Early", "Late"),
  method = "spearman",
  padjust = "fdr"
)

# Plotting results

# Load significant results from an RDS file (if available)
sign <- readRDS("fdr_sign.rds")
tail(sign)

# Write the significant results to an Excel file using openxlsx
library(openxlsx)
write.xlsx(sign, file = "genewise_DCA_results.xlsx")

# Define two genes of interest for plotting
geneA <- "Ywhab"
geneB <- "Map2k4"

# Create a data frame for plotting expression of the two genes across age groups
plotData <- data.frame(
  geneA = expr[geneA, ],
  geneB = expr[geneB, ],
  ageGroups = factor(samplesheet$Age_3bins, levels = c("Early", "Middle", "Late"))
)

# Calculate Spearman correlation between geneA and geneB in the 'Late' age group
correlation_late <- cor(
  plotData$geneA[plotData$ageGroups == "Late"],
  plotData$geneB[plotData$ageGroups == "Late"],
  method = "spearman"
)
print(paste("Spearman correlation in Late group:", correlation_late))

# Plot geneA vs geneB expression colored by age groups using ggplot2
library(ggplot2)
ggplot(plotData, aes(x = geneA, y = geneB, color = ageGroups)) +
  geom_point() +
  facet_wrap(~ ageGroups) +
  geom_smooth(method = "lm", se = FALSE) +
  labs(
    x = geneA,
    y = geneB,
    title = paste("Expression of", geneA, "vs", geneB, "by Age Group")
  ) +
  theme_minimal()

# Filter significant results from differential correlation analysis
significant_results <- subset(difcor_res, P <= 5e-5)
significant_results <- significant_results[!is.infinite(significant_results$zdiff), ] # Remove infinite zdiff
significant_results <- na.omit(significant_results)                                   # Remove NA values

# Load igraph library for network analysis
library(igraph)

# Create a network from the significant gene pairs
network <- makeNetwork(significant_results)

# Calculate degree centrality for each gene in the network
centrality <- getCentrality(network)

# Display the top 100 genes with highest degree centrality
head(centrality, 100)

# Display the distribution of node degrees
table(centrality$degree)

# Extract genes connected to 'Ywhab' and save them to a CSV file
psm <- significant_results[significant_results$GeneA == "Ywhab" | significant_results$GeneB == "Ywhab", ]
psm <- unique(c(psm$GeneA, psm$GeneB))
# Remove any suffix after '.' in gene names
psm <- sapply(strsplit(psm, split = "[.]"), function(x) x[1])
write.csv2(psm, file = "Ywhab_connected_genes.csv", quote = FALSE, row.names = FALSE)

# Display the first few rows of significant results
head(significant_results)

# Get hub enrichment with aging genes

# 'diffex_sign' is a vector of differentially expressed genes; ensure it's defined
# For example, load from a file or define based on analysis
# diffex_sign <- read.csv("diffex_sign.csv", stringsAsFactors = FALSE)$gene

# Identify top 5% of genes with highest degree (hubs)
top5percent <- centrality[1:round(0.05 * nrow(centrality)), ]
head(top5percent)

# Perform enrichment analysis to test if hubs are enriched for differentially expressed genes
enrich_res <- lapply(1:nrow(top5percent), function(i) {
  gene <- top5percent$genes[i]
  # Get genes connected to the hub gene
  psm <- significant_results[significant_results$GeneA == gene | significant_results$GeneB == gene, ]
  psm <- unique(c(psm$GeneA, psm$GeneB))
  psm <- psm[psm != gene]
  # Define background as all genes in the expression data
  background <- rownames(expr)
  # Ensure 'diffex_sign' is defined before running this
  # Create a contingency table for Fisher's exact test
  in_psm <- background %in% psm
  in_diffex <- background %in% diffex_sign
  contingency_table <- matrix(c(
    sum(in_psm & in_diffex),
    sum(in_psm & !in_diffex),
    sum(!in_psm & in_diffex),
    sum(!in_psm & !in_diffex)
  ), nrow = 2)
  res <- fisher.test(contingency_table)
  # Store results in a data frame
  out <- data.frame(gene = gene, p_value = res$p.value, odds_ratio = res$estimate)
  return(out)
})

# Combine enrichment results into a single data frame
enrich_res <- do.call(rbind, enrich_res)

# Sort enrichment results by p-value
enrich_res <- enrich_res[order(enrich_res$p_value), ]
head(enrich_res)

# Perform enrichment analysis for gene 'Gabrr2'
gene <- "Gabrr2"
psm <- significant_results[significant_results$GeneA == gene | significant_results$GeneB == gene, ]
psm <- unique(c(psm$GeneA, psm$GeneB))
psm <- psm[psm != gene]
background <- rownames(expr)
# Create a contingency table for Fisher's exact test
in_psm <- background %in% psm
in_diffex <- background %in% diffex_sign
contingency_table <- matrix(c(
  sum(in_psm & in_diffex),
  sum(in_psm & !in_diffex),
  sum(!in_psm & in_diffex),
  sum(!in_psm & !in_diffex)
), nrow = 2)
res <- fisher.test(contingency_table)
print(res)

# Get overlapping genes between psm and differentially expressed genes
overlapping_genes <- psm[psm %in% diffex_sign]
print(overlapping_genes)

# Find overlap between top hubs and differentially expressed genes
overlapping_hubs <- intersect(top5percent$genes, diffex_sign)
print(overlapping_hubs)

# Write centrality data to a CSV file
write.csv2(centrality, file = "initial_hubs.csv")

# Access centrality of gene 'Stat3'
centrality_stat3 <- centrality[centrality$genes == "Stat3", ]
print(centrality_stat3)

# Read pathway data from a CSV file
paths <- read.csv("./FEA_gProfiler_mmusculus_24-08-2022_09-24-24__intersections.csv", stringsAsFactors = FALSE)
```
