## Supplemental Appendix 1 for "Genetic modulation of immune gene co-expression in the aged mouse hippocampus by the *Apbb1ip* locus": utils_clean.R

```
# Function to flatten a correlation matrix into a data frame
flattenCorrMatrix <- function(cormat) {
  # Get the indices of the upper triangle of the correlation matrix
  upper_triangle <- upper.tri(cormat)
  
  # Create a data frame with row names, column names, and correlation values from the upper triangle
  data.frame(
    row = rownames(cormat)[row(cormat)[upper_triangle]],
    column = colnames(cormat)[col(cormat)[upper_triangle]],
    correlation = cormat[upper_triangle]
  )
}

# Function to compute the difference in Fisher's z-transformed correlations
diffCor <- function(rho1, n1, rho2, n2, method = "pearson") {
  # Apply Fisher's z-transformation to the correlation coefficients
  z1 <- atanh(rho1)
  z2 <- atanh(rho2)
  
  # Calculate the difference in z-transformed correlations
  z_diff <- z2 - z1
  
  # Compute the standard error based on the method
  if (method == "pearson") {
    se_diff <- sqrt((1 / (n1 - 3)) + (1 / (n2 - 3)))
  } else if (method == "spearman") {
    # Adjusted standard error for Spearman's correlation
    se_diff <- sqrt((1.06 / (n1 - 3)) + (1.06 / (n2 - 3)))
  } else {
    stop("Unsupported method. Use 'pearson' or 'spearman'.")
  }
  
  # Calculate the z-score difference
  z_score_diff <- z_diff / se_diff
  
  return(z_score_diff)
}

# Function to calculate two-tailed p-values from z-scores
getPvalue <- function(z_scores) {
  # Calculate two-tailed p-values from z-scores
  p_values <- 2 * pnorm(-abs(z_scores))
  return(p_values)
}

# Function to perform permutation for multiple testing correction (version 2)
permute_mtc2 <- function(data, samplesheet, genesA, genesB, contrastCol, contrast, method) {
  # Permute the labels in the contrast column
  samplesheet[, contrastCol] <- sample(samplesheet[, contrastCol])
  
  # Get the sample names for each group in the contrast
  group1_samples <- samplesheet[samplesheet[, contrastCol] == contrast[1], "colnames"]
  group2_samples <- samplesheet[samplesheet[, contrastCol] == contrast[2], "colnames"]
  
  # Calculate the correlation matrix between genesA and genesB for group 1
  ref_cor <- cor(
    t(data[genesA, group1_samples]),
    t(data[genesB, group1_samples]),
    method = method
  )
  
  # Calculate the correlation matrix between genesA and genesB for group 2
  alt_cor <- cor(
    t(data[genesA, group2_samples]),
    t(data[genesB, group2_samples]),
    method = method
  )
  
  # Convert the correlation matrices to data frames
  ref_cor_melted <- as.data.frame(as.table(ref_cor))
  alt_cor_melted <- as.data.frame(as.table(alt_cor))
  
  # Rename columns
  colnames(ref_cor_melted) <- c("GeneA", "GeneB", "ct_cor")
  ref_cor_melted$AD_cor <- alt_cor_melted$Freq
  
  # Get the sample sizes for each group
  ref_cor_melted$ct_N <- length(group1_samples)
  ref_cor_melted$ad_N <- length(group2_samples)
  
  # Calculate z-score differences between the correlations
  ref_cor_melted$zdiff <- diffCor(
    ref_cor_melted$ct_cor,
    ref_cor_melted$ct_N,
    ref_cor_melted$AD_cor,
    ref_cor_melted$ad_N,
    method = method
  )
  
  # Calculate p-values from z-score differences
  ref_cor_melted$P <- getPvalue(ref_cor_melted$zdiff)
  
  # Return the p-values
  return(ref_cor_melted$P)
}

# Function to perform permutation for multiple testing correction
permute_mtc <- function(data, samplesheet, contrastCol, contrast, method) {
  # Permute the labels in the contrast column
  samplesheet[, contrastCol] <- sample(samplesheet[, contrastCol])
  
  # Get sample names for each group
  group1_samples <- samplesheet[samplesheet[, contrastCol] == contrast[1], "colnames"]
  group2_samples <- samplesheet[samplesheet[, contrastCol] == contrast[2], "colnames"]
  
  # Calculate the correlation matrix for group 1
  ref_cor_matrix <- cor(
    t(data[, group1_samples]),
    method = method
  )
  
  # Calculate the correlation matrix for group 2
  alt_cor_matrix <- cor(
    t(data[, group2_samples]),
    method = method
  )
  
  # Flatten the correlation matrices into data frames
  ref_cor_flat <- flattenCorrMatrix(ref_cor_matrix)
  alt_cor_flat <- flattenCorrMatrix(alt_cor_matrix)
  
  # Rename columns
  colnames(ref_cor_flat) <- c("GeneA", "GeneB", "ct_cor")
  
  # Add the alternative group's correlations
  ref_cor_flat$AD_cor <- alt_cor_flat$correlation
  
  # Get sample sizes for each group
  ref_cor_flat$ct_N <- length(group1_samples)
  ref_cor_flat$ad_N <- length(group2_samples)
  
  # Calculate z-score differences
  ref_cor_flat$zdiff <- diffCor(
    ref_cor_flat$ct_cor,
    ref_cor_flat$ct_N,
    ref_cor_flat$AD_cor,
    ref_cor_flat$ad_N,
    method = method
  )
  
  # Calculate p-values
  ref_cor_flat$P <- getPvalue(ref_cor_flat$zdiff)
  
  # Return the p-values
  return(ref_cor_flat$P)
}

# Function to run differential correlation analysis
difcorRun <- function(data, samplesheet, contrastCol, contrast, method, padjust, Nperm = NULL) {
  # Get sample names for each group
  group1_samples <- samplesheet[samplesheet[, contrastCol] == contrast[1], "colnames"]
  group2_samples <- samplesheet[samplesheet[, contrastCol] == contrast[2], "colnames"]
  
  # Calculate correlation matrices for each group
  ref_cor_matrix <- cor(t(data[, group1_samples]), method = method)
  alt_cor_matrix <- cor(t(data[, group2_samples]), method = method)
  
  # Flatten correlation matrices into data frames
  ref_cor_flat <- flattenCorrMatrix(ref_cor_matrix)
  alt_cor_flat <- flattenCorrMatrix(alt_cor_matrix)
  
  # Rename columns
  colnames(ref_cor_flat) <- c("GeneA", "GeneB", "ct_cor")
  
  # Add the alternative group's correlations
  ref_cor_flat$AD_cor <- alt_cor_flat$correlation
  
  # Get sample sizes for each group
  ref_cor_flat$ct_N <- length(group1_samples)
  ref_cor_flat$ad_N <- length(group2_samples)
  
  # Calculate z-score differences
  ref_cor_flat$zdiff <- diffCor(
    ref_cor_flat$ct_cor,
    ref_cor_flat$ct_N,
    ref_cor_flat$AD_cor,
    ref_cor_flat$ad_N,
    method = method
  )
  
  # Calculate p-values
  ref_cor_flat$P <- getPvalue(ref_cor_flat$zdiff)
  
  # Adjust p-values based on the specified method
  if (padjust == "fdr") {
    # Adjust p-values using false discovery rate
    ref_cor_flat$fdr <- p.adjust(ref_cor_flat$P, method = "fdr")
    ref_cor_flat <- ref_cor_flat[order(ref_cor_flat$fdr), ]
  } else if (padjust == "perm") {
    # Adjust p-values using permutation test
    ref_cor_flat <- ref_cor_flat[order(ref_cor_flat$P), ]
    
    nominal_pvalues <- ref_cor_flat$P
    perm_corrected <- vector(mode = "integer", length = length(nominal_pvalues))
    
    # Perform permutations
    for (i in seq_len(Nperm)) {
      if (i %% 10 == 0) {
        message(sprintf("%d of %d permutations done...", i, Nperm))
      }
      # Get permuted p-values
      permuted_pvalues <- permute_mtc(data, samplesheet, contrastCol, contrast, method)
      permuted_pvalues <- sort(permuted_pvalues)
      
      # Count how many permuted p-values are less than nominal p-values
      perm_corrected <- perm_corrected + findInterval(nominal_pvalues, permuted_pvalues)
    }
    
    # Calculate permutation-based p-values
    perm_corrected <- perm_corrected / (length(nominal_pvalues) * Nperm)
    ref_cor_flat$perm_P <- perm_corrected
  }
  
  return(ref_cor_flat)
}

# Function to create a network from the results
makeNetwork <- function(results) {
  # Load necessary library
  require(igraph)
  
  # Create a data frame of edges (links) from results
  edges <- data.frame(
    source = as.character(results$GeneA),
    target = as.character(results$GeneB)
  )
  
  # Get the unique list of genes (nodes)
  genes <- unique(c(edges$source, edges$target))
  nodes <- data.frame(name = genes)
  
  # Create an undirected graph from the data frames
  network <- graph_from_data_frame(d = edges, vertices = nodes, directed = FALSE)
  
  return(network)
}

# Function to calculate degree centrality of the network nodes
getCentrality <- function(network) {
  # Calculate the degree centrality of each node in the network
  degree_centrality <- degree(network)
  
  # Convert to a data frame
  centrality_df <- data.frame(
    genes = names(degree_centrality),
    degree = degree_centrality
  )
  
  # Order by degree descending
  centrality_df <- centrality_df[order(centrality_df$degree, decreasing = TRUE), ]
  
  # Add an index column
  centrality_df$index <- seq_len(nrow(centrality_df))
  
  return(centrality_df)
}
```
