## Supplemental Appendix 1 for "Genetic modulation of immune gene co-expression in the aged mouse hippocampus by the *Apbb1ip* locus": WGCNA_GOanalysis_GeneOverlap.R

```
####

inputDir             <- workingDir
outputDir            <- "/Users/methiaa/Downloads/GroupC_BulkRNA_WGCNA/output/"
normCountsFile       <- "normalized_counts.csv"
traitDataFilePath    <- "/Users/methiaa/Downloads/GroupC_BulkRNA_WGCNA/trait_data.csv"
projectName          <- "GroupC_WGCNA_binned"

cat("1) Loading WGCNA library.. \n")
# Load the WGCNA package
suppressPackageStartupMessages(library("WGCNA", warn.conflicts=FALSE, quietly=TRUE))
# The following setting is important, do not omit.
options(stringsAsFactors = FALSE);

# DATA INPUT AND PREPROCESSING

cat("2) Reading in the input gene data set.. \n")
#Read in the normalized gene count data set
geneData = readRDS(paste(inputDir, "normalized_counts.rds", sep = '' ));
# geneData = read.csv(paste(inputDir, normCountsFile, sep = '' ), header = F);
# colnames(geneData) <- geneData[1,]
# geneData <- geneData[-1,]

# Take a quick look at what is in the data set:
print(dim(geneData))
print(names(geneData))

#rownames(geneData) <- geneData[,1]
# datExpr0 = as.data.frame(t(geneData[, -c(1)]));
datExpr0 = as.data.frame(t(geneData));
# names(datExpr0) = geneData$X;
# rownames(datExpr0) = names(geneData)[-c(1)];

gsg = goodSamplesGenes(datExpr0, verbose = 3);
print(gsg$allOK)

if (!gsg$allOK)
{
  # Optionally, print the gene and sample names that were removed:
  if (sum(!gsg$goodGenes)>0) 
    printFlush(paste("Removing genes:", paste(names(datExpr0)[!gsg$goodGenes], collapse = ", ")));
  if (sum(!gsg$goodSamples)>0) 
    printFlush(paste("Removing samples:", paste(rownames(datExpr0)[!gsg$goodSamples], collapse = ", ")));
  # Remove the offending genes and samples from the data:
  datExpr0 = datExpr0[gsg$goodSamples, gsg$goodGenes]
}

# SAMPLE CLUSTERING

cat("3) Clustering samples.. \n")

sampleTree = hclust(dist(datExpr0), method = "average");
# Plot the sample tree: Open a graphic output window of size 12 by 9 inches
# The user should change the dimensions if the window is too large or too small.
# sizeGrWindow(12,9)
pdf(file = paste(outputDir,"sampleClustering.pdf", sep = '' ), width = 20, height = 8);
par(cex = 0.6);
par(mar = c(0,4,2,0))
plot(sampleTree, main = "Sample clustering to detect outliers", sub="", xlab="", cex.lab = 1.5, 
     cex.axis = 1.5, cex.main = 2)
# Plot a line to show the cut
abline(h = 52, col = "red");
dev.off()

# If there are any sample outliers:

# Determine cluster under the line
clust = cutreeStatic(sampleTree, cutHeight = 52, minSize = 10)
table(clust)
# clust 1 contains the samples we want to keep.
keepSamples = (clust==1)
datExpr = datExpr0[keepSamples, ]
mismatched.samples <- c("BXD65b_1","BXD75_1")
datExpr <- datExpr[!(rownames(datExpr) %in% mismatched.samples), ]

#datExpr = datExpr0
nGenes = ncol(datExpr)
nSamples = nrow(datExpr)
#sampleIDs <- substr(rownames(datExpr),2,5)
sampleIDs <- rownames(datExpr)
rownames(datExpr) <- sampleIDs

# LOAD BEHAVIORAL/PHENO. TRAIT DATA

cat("4) Reading in the trait data set.. \n")

suppressPackageStartupMessages(library("dplyr", warn.conflicts=FALSE, quietly=TRUE))
suppressPackageStartupMessages(library("tidyr", warn.conflicts=FALSE, quietly=TRUE))

phenoData = read.csv(traitDataFilePath, header = T);
print(dim(phenoData))
print(names(phenoData))
names(phenoData)[1] <- "Sample"

reqdSamples = rownames(datExpr);
traitRows = match(reqdSamples, phenoData$Sample);

datTraits = phenoData[traitRows, ];
rownames(datTraits) = phenoData[traitRows, 1];

phenoData <- phenoData %>% 
  mutate(var = 1) %>% 
  spread(Age_3bins, var, fill = 0) %>% 
  left_join(phenoData) %>% 
  select(Sample, Age, Age_3bins, everything())

# Form a data frame analogous to expression data that will hold the clinical traits.

reqdSamples = rownames(datExpr);
traitRows = match(reqdSamples, phenoData$Sample);

datTraits = phenoData[traitRows, -c(1,3)];
rownames(datTraits) = phenoData[traitRows, 1];

### Gene expression heatmap ----

# suppressPackageStartupMessages(library("pheatmap", warn.conflicts=FALSE, quietly=TRUE))
# suppressPackageStartupMessages(library("RColorBrewer", warn.conflicts=FALSE, quietly=TRUE))
# 
# select1 <- phenoData[traitRows,][, c("Sample", "Age_3bins")];
# select1$Age_3bins <- factor(select1$Age_3bins)
# 
# datExpr_t <- t(datExpr)
# 
# annot_col <- as.data.frame(select1[2])
# row.names(annot_col) <- select1$Sample
# 
# traitColors1 <- labels2colors(levels(annot_col$Age_3bins), colorSeq = brewer.pal(n = length(levels(annot_col$Age_3bins)), name = "BrBG"))
# names(traitColors1) <- levels(annot_col$Age_3bins)
# traitColors <- list(traitColors1)
# names(traitColors) <- c("Age_3bins")
# 
# select_genes <- order(rowMeans(datExpr_t),
#                       decreasing=TRUE)[1:nGenes]
# counts.heatmap <- datExpr_t[select_genes,]
# 
# # counts.heatmap <- t(scale(x=t(counts.heatmap), center=TRUE))
# 
# p <- pheatmap(counts.heatmap,
#          show_rownames=FALSE,
#          cluster_cols=TRUE,
#          treeheight_row = 0,
#          # cluster_rows = FALSE,
#          scale = "row",
#          annotation_col = annot_col,
#          annotation_names_col = FALSE,
#          color = colorRampPalette(rev(brewer.pal(n = 7, name = "RdBu")))(100),
#          #cutree_rows = 2,
#          #annotation_names_col = FALSE,
#          angle_col = 90,
#          annotation_colors = traitColors,
#          show_colnames = TRUE)
# pdf(file = paste(outputDir, "heatmap_annot.pdf", sep = ''), width = 6, height = 5);
# print(p)
# dev.off()

####

collectGarbage();

#### Re-cluster samples along with traits: ----

cat("5) Re-clustering the samples with traits.. \n")

sampleTree2 = hclust(dist(datExpr), method = "average")

# Convert traits to a color representation: white means low, red means high, grey means missing entry
#traitColors = numbers2colors(datTraits, signed = FALSE);

traitColors <- data.frame(matrix(ncol=length(names(datTraits)),nrow=nSamples, dimnames=list(NULL, names(datTraits))))
#n = length(levels(as.factor(datTraits[,trait])))
for (trait in names(datTraits)){
  #select <- phenoData[traitRows,][, c("Sample", trait)];
  traitColors[, trait] = numbers2colors(datTraits[,trait], 
                                        signed = FALSE,
                                        #colors = brewer.pal(n = length(levels(as.factor(datTraits[,trait]))), name = "Set3"),
                                        #colors = blueWhiteRed(4)[3:4],
                                        centered = TRUE)
}

# Plot the sample dendrogram and the colors underneath.
pdf(file = paste(outputDir,"sampleClustering_traits.pdf", sep = '' ), width = 20, height = 6);
plotDendroAndColors(sampleTree2, traitColors,
                    groupLabels = names(datTraits), 
                    cex.dendroLabels = 0.5, 
                    autoColorHeight = FALSE, 
                    colorHeight = 0.2,
                    main = "Sample dendrogram and trait heatmap")
dev.off()

save(datExpr, datTraits, file = paste(outputDir, sprintf("R_Data/%s-dataInput.RData", projectName), sep = ''))

cat("6) Starting network topology analysis.. \n")

# Choose a set of soft-thresholding powers
powers = c(c(1:10), seq(from = 12, to=30, by=2))
# Call the network topology analysis function
sft = pickSoftThreshold(datExpr, powerVector = powers, networkType = "signed", verbose = 5)

write.table(as.data.frame(sft), file = paste(outputDir, sprintf("/%s-sft_table.txt", projectName), sep = ''), quote = FALSE)

# Plot the results:
# sizeGrWindow(9, 5)
pdf(file = paste(outputDir,"softThesholding_connectivity_plot.pdf", sep = '' ), width = 8.5, height = 4.5);
par(mfrow = c(1,2));
cex1 = 0.7;
# Scale-free topology fit index as a function of the soft-thresholding power
plot(sft$fitIndices[,1], -sign(sft$fitIndices[,3])*sft$fitIndices[,2],
     xlab="Soft Threshold (power)",ylab="Scale Free Topology Model Fit,signed R^2",type="n",
     main = paste("Scale independence"));
text(sft$fitIndices[,1], -sign(sft$fitIndices[,3])*sft$fitIndices[,2],
     labels=powers,cex=cex1,col="red");
# this line corresponds to using an R^2 cut-off of h
abline(h=0.9,col="red")
# Mean connectivity as a function of the soft-thresholding power
plot(sft$fitIndices[,1], sft$fitIndices[,5],
     xlab="Soft Threshold (power)",ylab="Mean Connectivity", type="n",
     main = paste("Mean connectivity"))
text(sft$fitIndices[,1], sft$fitIndices[,5], labels=powers, cex=cex1,col="red")
dev.off()

## After selecting soft power from the table above 

soft_power       <- 10
outputDir        <- "/Users/methiaa/Downloads/GroupC_BulkRNA_WGCNA/output/"
projectName      <- "GroupC_WGCNA_binned"

# # Load the WGCNA package
# suppressPackageStartupMessages(library("WGCNA", warn.conflicts=FALSE, quietly=TRUE))
library(gprofiler2)

# The following setting is important, do not omit.
options(stringsAsFactors = FALSE);

# Load the expression and trait data saved in the first part
lnames = load(file = paste(outputDir, sprintf("R_Data/%s-dataInput.RData", projectName), sep = ''));
#The variable lnames contains the names of loaded variables.
lnames

###### Alternative: one step blockwise modules ----

workingDir = "/Users/methiaa/Downloads/GroupC_BulkRNA_WGCNA/output/R_Data/";
setwd(workingDir);

bwnet = blockwiseModules(datExpr, maxBlockSize = 2000,
                         power = 10, TOMType = "signed", minModuleSize = 30,
                         reassignThreshold = 0, mergeCutHeight = 0.2,
                         numericLabels = TRUE,
                         deepSplit = 3,
                         pamRespectsDendro = FALSE,
                         loadTOM = TRUE,
                         #saveTOMs = TRUE,
                         #saveTOMFileBase = "GroupC_wgcna",
                         verbose = 3)

# # open a graphics window
# sizeGrWindow(12, 9)
# # Convert labels to colors for plotting
# mergedColors = labels2colors(bwnet$colors)
# # Plot the dendrogram and the module colors underneath
# plotDendroAndColors(bwnet$dendrograms[[1]], mergedColors[bwnet$blockGenes[[1]]],
#                     "Module colors",
#                     dendroLabels = FALSE, hang = 0.03,
#                     addGuide = TRUE, guideHang = 0.05)

moduleLabels = bwnet$colors
moduleColors = labels2colors(bwnet$colors)
# MEs = bwnet$MEs;
# geneTree = bwnet$dendrograms[[1]];

# Calculate eigengenes
MEList = moduleEigengenes(datExpr, colors = moduleColors)
MEs = MEList$eigengenes
# Calculate dissimilarity of module eigengenes
MEDiss = 1-cor(MEs);
# Cluster module eigengenes
METree = hclust(as.dist(MEDiss), method = "average");
MEDissThres = 0.2

# Plot the result
pdf(file = paste(outputDir,"ME_clustering_bwnet.pdf", sep = '' ), width = 7, height = 6);
plot(METree, main = NULL, cex = 0.8,
     xlab = "", sub = "")
# # Plot the cut line into the dendrogram
# abline(h=MEDissThres, col = "red")
dev.off()

#######

cat("7) Calculating adjacency matrix.. \n")

softPower = soft_power;
adjacency = adjacency(datExpr, power = softPower);

cat("8) Calculating TOM and dissimilarity matrix.. \n")

# Turn adjacency into topological overlap
TOM = TOMsimilarity(adjacency);
dissTOM = 1-TOM

cat("9) Identifying modules.. \n")

# Call the hierarchical clustering function
geneTree = hclust(as.dist(dissTOM), method = "average");
# Plot the resulting clustering tree (dendrogram)
# sizeGrWindow(12,9)
# plot(geneTree, xlab="", sub="", main = "Gene clustering on TOM-based dissimilarity",
#      labels = FALSE, hang = 0.04);

# We like large modules, so we set the minimum module size relatively high:
minModuleSize = 30;
# Module identification using dynamic tree cut:
dynamicMods = cutreeDynamic(dendro = geneTree, distM = dissTOM,
                            deepSplit = 3, pamRespectsDendro = FALSE,
                            minClusterSize = minModuleSize);
table(dynamicMods)

# Convert numeric lables into colors
dynamicColors = labels2colors(dynamicMods)
table(dynamicColors)
# Plot the dendrogram and colors underneath
# sizeGrWindow(8,6)
# pdf(file = paste(outputDir,"dendrogram_modules.pdf", sep = '' ), width = 9, height = 6);
# plotDendroAndColors(geneTree, dynamicColors, "Dynamic Tree Cut",
#                     dendroLabels = FALSE, hang = 0.03,
#                     addGuide = TRUE, guideHang = 0.05,
#                     main = "Gene dendrogram and module colors")
# dev.off()

cat("10) Calculating and clustering module eigengenes (MEs).. \n")

# Calculate eigengenes
MEList = moduleEigengenes(datExpr, colors = dynamicColors)
MEs = MEList$eigengenes
# Calculate dissimilarity of module eigengenes
MEDiss = 1-cor(MEs);
# Cluster module eigengenes
METree = hclust(as.dist(MEDiss), method = "average");
MEDissThres = 0.2

# Plot the result
sizeGrWindow(7, 6)
pdf(file = paste(outputDir,"ME_clustering.pdf", sep = '' ), width = 7, height = 7);
plot(METree, main = "Clustering of module eigengenes",
     xlab = "", sub = "")
# Plot the cut line into the dendrogram
abline(h=MEDissThres, col = "red")
dev.off()

# Call an automatic merging function
merge = mergeCloseModules(datExpr, dynamicColors, cutHeight = MEDissThres, verbose = 3)
# The merged module colors
mergedColors = merge$colors;

table(mergedColors)

# Eigengenes of the new merged modules:
mergedMEs = merge$newMEs;

cat("11) Plotting gene dendrogram with merged modules.. \n")

# sizeGrWindow(12, 9)
pdf(file = paste(outputDir,"geneDendro_merged.pdf", sep = '' ), width = 9, height = 6)
plotDendroAndColors(geneTree, cbind(dynamicColors, mergedColors),
                    c("Dynamic Tree Cut", "Merged dynamic"),
                    dendroLabels = FALSE, hang = 0.03,
                    addGuide = TRUE, guideHang = 0.05)
dev.off()


# Rename to moduleColors
moduleColors = mergedColors
# Construct numerical labels corresponding to the colors
colorOrder = c("grey", standardColors(50));
moduleLabels = match(moduleColors, colorOrder)-1;
MEs = mergedMEs;
# Save module colors and labels for use in subsequent parts
save(TOM, dissTOM, MEs, moduleLabels, moduleColors, geneTree, file = paste(outputDir, sprintf("R_Data/%s-networkConstruction-stepByStep.RData", projectName), sep=""))

# load(file = paste(outputDir, sprintf("R_Data/%s-networkConstruction-stepByStep.RData", projectName), sep=""))

###########

nGenes = ncol(datExpr)
nSamples = nrow(datExpr)

cat("12) Recalculating MEs with color labels.. \n")

# Recalculate MEs with color labels
MEs0 = moduleEigengenes(datExpr, moduleColors)$eigengenes
MEs = orderMEs(MEs0)
moduleTraitCor = cor(MEs, datTraits, use = "p");
moduleTraitPvalue = corPvalueStudent(moduleTraitCor, nSamples);

# Will display correlations and their p-values
textMatrix =  paste(signif(moduleTraitCor, 2), "\n(",
                    signif(moduleTraitPvalue, 1), ")", sep = "");
dim(textMatrix) = dim(moduleTraitCor)

cat("13) Plotting module-trait relationships (correlations and p-values).. \n")

# Display the correlation values within a heatmap plot
pdf(file = paste(outputDir,"module_trait_relationships_ds3.pdf", sep = '' ), width = 5, height = 14);
par(mar = c(6, 9, 3, 3));
labeledHeatmap(Matrix = moduleTraitCor,
               xLabels = names(datTraits),
               yLabels = names(MEs),
               ySymbols = names(MEs),
               colorLabels = FALSE,
               colors = blueWhiteRed(50),
               textMatrix = textMatrix,
               setStdMargins = FALSE,
               cex.text = 0.75,
               zlim = c(-1,1),
               main = paste("Module-trait relationships"))
dev.off()

pdf(file = paste(outputDir,"module_trait_relationships_filtered.pdf", sep = '' ), width = 5, height = 2.9);
par(mar = c(6, 9, 3, 3));
# labeledHeatmap(Matrix = moduleTraitCor[c(9,21,26,29),c(1,2,4,3)],
#                xLabels = names(datTraits)[c(1,2,4,3)],
#                yLabels = names(MEs)[c(9,21,26,29)],
#                ySymbols = names(MEs)[c(9,21,26,29)],
#                colorLabels = FALSE,
#                colors = blueWhiteRed(50),
#                textMatrix = textMatrix[c(9,21,26,29),c(1,2,4,3)],
#                setStdMargins = FALSE,
#                cex.text = 0.8,
#                zlim = c(-1,1),
#                main = paste("Module-trait relationships"))
labeledHeatmap(Matrix = moduleTraitCor[c(9,18),c(1,2,4,3)],
               xLabels = names(datTraits)[c(1,2,4,3)],
               yLabels = names(MEs)[c(9,18)],
               ySymbols = names(MEs)[c(9,18)],
               colorLabels = FALSE,
               #colors = brewer.pal(10, "PuOr"),
               colors = blueWhiteRed(50),
               textMatrix = textMatrix[c(9,18),c(1,2,4,3)],
               setStdMargins = FALSE,
               cex.text = 0.8,
               zlim = c(-1,1),
               main = paste("Module-trait relationships"))
dev.off()

# names (colors) of the modules
modNames = substring(names(MEs), 3)

## proportion of variance explained by eigengenes
pvEX <- propVarExplained(datExpr, moduleColors, MEs, corFnc = "cor", corOptions = "use = 'p'")
write.csv(as.data.frame(pvEX), file = paste(outputDir, "propVarExplained.csv", sep = ""), quote = F)

cat("14) Calculating module membership for all genes.. \n")

geneModuleMembership = as.data.frame(cor(datExpr, MEs, use = "p"));
MMPvalue = as.data.frame(corPvalueStudent(as.matrix(geneModuleMembership), nSamples));
names(geneModuleMembership) = paste("MM", modNames, sep="");
names(MMPvalue) = paste("p.MM", modNames, sep="");

cat("15) Running calculations for all traits.. \n ")

for (trait in names(datTraits)){
  
  #output_dir       <- dirname(outputDir);
  trait_resultsDir      <- paste(outputDir, trait, "/", sep='')
  system(paste("mkdir -p", trait_resultsDir, sep=' '))
  
  cat(sprintf("Running calculations for the trait: '%s'.. \n", trait))
  
  t <- as.data.frame(moduleTraitCor, row.names = row.names(moduleTraitCor))
  q <- as.data.frame(moduleTraitPvalue, row.names = row.names(moduleTraitPvalue))
  t[, paste(trait, "p_value", sep = '_')] <- q[, trait]
  sel_modules <- row.names(t)[(t[, trait]>0.15 | t[, trait]<(-0.15)) & t[, paste(trait, "p_value", sep = '_')]<0.05]
  num_modules <- length(sel_modules)
  sel_modules = substring(sel_modules, 3)
  
  # Define variable <trait> containing that trait column of datTrait
  score_trait = as.data.frame(datTraits[, trait]);
  names(score_trait) = trait
  
  geneTraitSignificance = as.data.frame(cor(datExpr, score_trait, use = "p"));
  GSPvalue = as.data.frame(corPvalueStudent(as.matrix(geneTraitSignificance), nSamples));
  names(geneTraitSignificance) = paste("GS.", names(score_trait), sep="");
  names(GSPvalue) = paste("p.GS.", names(score_trait), sep="");
  
  if (num_modules != 0) {
    
    moduleGenes <- data.frame(Logicals=logical(nGenes),
                              stringsAsFactors=F)
    moduleGenes <- moduleGenes[rep(names(moduleGenes), c(num_modules))]
    column <- vector(mode = "integer", length = num_modules)
    # module_vec <- vector(mode = "character", length = num_modules)
    
    for (i in 1:num_modules)
    {
      module = sel_modules[i]
      column[i] = match(module, modNames);
      moduleGenes_sel = moduleColors==module;
      moduleGenes[1:nGenes,i] <- moduleGenes_sel
    }
    
    cat(sprintf("\t Plotting module membership vs. gene significance for the trait: '%s'.. \n", trait))
    
    # pdf(file = paste(outputDir,"MM_GS_CS_all.pdf", sep = '')); # ), height = 5*(num_modules_all %/% 2), width = 5*(num_modules_all %/% (num_modules_all %/% 2))+1);
    # if (num_modules_all %/% 2 != 0) {
    #   par(mfrow = c(num_modules_all %/% 2, (num_modules_all %/% (num_modules_all %/% 2))+1));
    # } else {
    #   par(mfrow = c(1, 1));
    # }
    for (i in 1:num_modules)
    {
      pdf(file = paste(trait_resultsDir, sel_modules[i],"_MM_GS_", trait, ".pdf", sep = ''), width = 5, height = 5);
      verboseScatterplot(abs(geneModuleMembership[moduleGenes[,i], column[i]]),
                         abs(geneTraitSignificance[moduleGenes[,i], 1]),
                         xlab = paste("Module Membership in", sel_modules[i], "module"),
                         ylab = paste("Gene significance for ", trait, sep = ''),
                         main = paste("Module membership vs. gene significance\n"),
                         cex.main = 1.2, cex.lab = 1.2, cex.axis = 1.2, col = sel_modules[i])
      dev.off()
      
      test_df <- data.frame(x= abs(geneModuleMembership[moduleGenes[,i], column[i]]),y=abs(geneTraitSignificance[moduleGenes[,i], 1]))
      sc_plot <- ggscatter(test_df, x = "x", y = "y",
                           add = "reg.line",  # Add regression line
                           add.params = list(color = "black", fill = "gray"), # Customize reg. line
                           cor.coef = TRUE,
                           cor.coeff.args = list(method = "pearson",
                                                 #label.x.npc = 1,
                                                 label.y.npc = 1,
                                                 label.x = 0.7),
                           color = sel_modules[i],
                           size = 1.75,
                           xlab = paste("Module Membership in", sel_modules[i], "module"),
                           ylab = paste("Gene significance for ", trait, sep = ''),
                           main = paste("Module membership vs. gene significance\n"),
                           font.label = c(11, "bold"),
                           conf.int = TRUE) + theme(panel.border = element_rect(colour = "black", fill=NA, size=1))
      pdf(file = paste(trait_resultsDir, sel_modules[i],"_MM_GS_final_", trait, ".pdf", sep = ''), width = 4, height = 4);
      print(sc_plot)
      dev.off()
    }
    
    # names(datExpr)
    # names(datExpr)[moduleColors=="purple"]
    
    # # Read in the annotation file
    # annot = read.csv(file = "GeneAnnotation.csv");
    # dim(annot)
    # names(annot)
    # probes = names(datExpr)
    # probes2annot = match(probes, annot$substanceBXH)
    # # The following is the number or probes without annotation:
    # sum(is.na(probes2annot))
    # # Should return 0.
    
    
    #cat("17) Writing results to CSV file.. \n")
    cat(sprintf("\t Writing results to CSV file for the trait: '%s'.. \n", trait))
    #cat("17.1) Writing results ordered by trait: CS_all.. \n")
    
    # Create the starting data frame
    geneInfo0 = data.frame(GeneIDs = names(datExpr),
                           #geneSymbol = annot$gene_symbol[probes2annot],
                           #LocusLinkID = annot$LocusLinkID[probes2annot],
                           moduleColor = moduleColors,
                           geneTraitSignificance,
                           GSPvalue)
    
    # Order modules by their significance for trait (eg. CS_all)
    modOrder = order(-abs(cor(MEs, score_trait, use = "p")));
    # Add module membership information in the chosen order
    for (mod in 1:ncol(geneModuleMembership))
    {
      oldNames = names(geneInfo0)
      geneInfo0 = data.frame(geneInfo0, geneModuleMembership[, modOrder[mod]], 
                             MMPvalue[, modOrder[mod]]);
      names(geneInfo0) = c(oldNames, paste("MM.", modNames[modOrder[mod]], sep=""),
                           paste("p.MM.", modNames[modOrder[mod]], sep=""))
    }
    # Order the genes in the geneInfo variable first by module color, then by geneTraitSignificance
    geneOrder = order(geneInfo0$moduleColor, -abs(geneInfo0[, paste("GS.", trait, sep = '')]));
    geneInfo = geneInfo0[geneOrder, ]
    
    # library(gprofiler2)
    annot_names <- gconvert(query = geneInfo$GeneIDs,
                            organism = "mmusculus",
                            target="ENSG",
                            mthreshold = 1, filter_na = FALSE)
    
    # library(biomaRt)
    # ensembl <- useMart("ensembl")
    # ensembl <- useDataset("mmusculus_gene_ensembl", mart = ensembl)
    # annot_names <- getBM(attributes = c("ensembl_gene_id", "external_gene_name"), 
    #                      filters = "ensembl_gene_id", 
    #                      values = geneInfo$EnsemblIDs, 
    #                      mart = ensembl,
    #                      uniqueRows = TRUE)
    # geneInfo = left_join(x = geneInfo, y = annot_names, by = c("EnsemblIDs" = "ensembl_gene_id"))
    # geneInfo$external_gene_name[is.na(geneInfo$external_gene_name)] <- "None"
    
    geneInfo["EnsemblIDs"] <- annot_names$target
    ncols_df <- length(names(geneInfo))
    geneInfo <- geneInfo[,c(1, ncols_df, 2:(ncols_df-1))]
    
    write.csv(geneInfo, file = paste(trait_resultsDir, projectName, "_trait_GSordered_allmod_geneInfo.csv", sep = ''))
    
    for (i in 1:num_modules)
    {
      # Select modules
      modules = sel_modules[i];
      
      # Create the starting data frame for each module
      modgeneInfo <- geneInfo[geneInfo$moduleColor == modules,]
      filtMM_modgeneInfo <- modgeneInfo[abs(modgeneInfo[, paste("MM.", modules, sep = '')])>0.5
                                        & modgeneInfo[, paste("p.MM.", modules, sep = '')]<0.05, ]
      
      filteredmodgenes <- modgeneInfo[abs(modgeneInfo[, paste("GS.", trait, sep = '')])>0.15
                                      & modgeneInfo[, paste("p.GS.", trait, sep = '')]<0.05
                                      & abs(modgeneInfo[, paste("MM.", modules, sep = '')])>0.5
                                      & modgeneInfo[, paste("p.MM.", modules, sep = '')]<0.05, ]
      
      filteredhubgenes <- modgeneInfo[abs(modgeneInfo[, paste("GS.", trait, sep = '')])>0.2 
                                      & modgeneInfo[, paste("p.GS.", trait, sep = '')]<0.05 
                                      & abs(modgeneInfo[, paste("MM.", modules, sep = '')])>0.8 
                                      & modgeneInfo[, paste("p.MM.", modules, sep = '')]<0.05, ]
      
      # cat(sprintf("\t Hub genes for the module '%s' and trait '%s'- %s: .. \n", modules, trait, filteredhubgenes$EnsemblIDs, filteredhubgenes$EntrezGeneSymbol))
      # cat(sprintf("\t Hub genes for the module '%s' and trait '%s': \n", modules, trait))
      # cat(sprintf("\t\t %s\t%s \n", filteredhubgenes$EnsemblIDs, filteredhubgenes$`Entrez Gene Symbol`))
      
      write.csv(filteredhubgenes, file = paste(trait_resultsDir, projectName, "_", modules, "_", "hubgeneInfo.csv", sep = ''))
      write.csv(filteredmodgenes, file = paste(trait_resultsDir, projectName, "_", modules, "_", "filteredgeneInfo.csv", sep = ''))
      write.csv(modgeneInfo, file = paste(trait_resultsDir, projectName, "_", modules, "_", "allgeneInfo.csv", sep = ''))
      #write.csv(filtMM_modgeneInfo, file = paste(trait_resultsDir, projectName, "_", modules, "_", "filtMMgeneInfo.csv", sep = ''))
      
      # Select module probes
      # probes = names(datExpr)
      # inModule = is.finite(match(moduleColors, modules));
      # modProbes = probes[inModule];
      filteredmodgenes <- filteredmodgenes %>% filter(EnsemblIDs != "None")
      inModule = match(filteredmodgenes[,"GeneIDs"],names(datExpr))
      modProbes = filteredmodgenes[,"GeneIDs"]
      # modGenes = annot$gene_symbol[match(modProbes, annot$substanceBXH)];
      
      if (length(modProbes) > 1){
        # Select the corresponding Topological Overlap
        modTOM = TOM[inModule, inModule];
        dimnames(modTOM) = list(modProbes, modProbes)
        
        # cat("21) Exporting Cytoscape input files.. \n")
        
        # Export the network into edge and node list files Cytoscape can read
        cyt = exportNetworkToCytoscape(modTOM,
                                       edgeFile = sprintf(paste(trait_resultsDir, paste("%s-CytoscapeInput-edges-", modules, ".txt", sep=""), sep=""), projectName),
                                       nodeFile = sprintf(paste(trait_resultsDir, paste("%s-CytoscapeInput-nodes-", modules, ".txt", sep=""), sep=""), projectName),
                                       weighted = TRUE,
                                       threshold = 0.00,
                                       nodeNames = modProbes,
                                       # altNodeNames = modGenes,
                                       nodeAttr = moduleColors[inModule])
        cyt = exportNetworkToCytoscape(modTOM,
                                       edgeFile = sprintf(paste(trait_resultsDir, paste("%s-CytoscapeInput-edges-filt-", modules, ".txt", sep=""), sep=""), projectName),
                                       nodeFile = sprintf(paste(trait_resultsDir, paste("%s-CytoscapeInput-nodes-filt-", modules, ".txt", sep=""), sep=""), projectName),
                                       weighted = TRUE,
                                       threshold = 0.05,
                                       nodeNames = modProbes,
                                       # altNodeNames = modGenes,
                                       nodeAttr = moduleColors[inModule])
      }
    }
  } else {
    print(paste("No significantly correlated modules found for trait: ", trait, sep = ''))
  }
}

saveRDS(geneInfo0, file = paste(outputDir,"geneInfo_allModules.rds", sep = '' ))
saveRDS(geneModuleMembership, file = paste(outputDir,"geneModuleMembership_allModules.rds", sep = '' ))
saveRDS(MEs0, file = paste(outputDir,"moduleEigengeneVectors_allModules.rds", sep = '' ))
write.csv(MEs, file = paste(outputDir,"moduleEigengeneVectors_allModules.csv", sep = '' ), quote = FALSE, row.names = TRUE)
write.csv(geneInfo0, file = paste(outputDir,"geneInfo_allModules.csv", sep = '' ), quote = FALSE, row.names = TRUE)
write.csv(geneModuleMembership, file = paste(outputDir,"geneModuleMembership_allModules.csv", sep = '' ), quote = FALSE, row.names = TRUE)
write.csv(geneInfo0$GeneIDs, file = paste(outputDir,"backgroundGenes.txt", sep = '' ), quote = FALSE, row.names = FALSE)

save(bwnet, MEs, moduleLabels, moduleColors, 
     file = paste(outputDir, sprintf("R_Data/%s-networkConstruction-outputvariables_blockwise.RData", projectName), sep=""))

# # Transform dissTOM with a power to make moderately strong connections more visible in the heatmap
# plotTOM = dissTOM^7;
# # Set diagonal to NA for a nicer plot
# diag(plotTOM) = NA;

# save(plotTOM, file = sprintf("/fischerlab/users/methia/IIDcogProject/rnaseq/WGCNA/R_Data/%s/%s-plotTOM.RData", brainRegion, brainRegion))

# cat("18.1) Plotting network heatmap plot with all genes.. \n")

# Call the plot function
# sizeGrWindow(9,9)
# pdf(file = paste(outputDir,"NetworkHeatmap_all_genes.pdf", sep = '' ), width = 9, height = 9);
# TOMplot(plotTOM, geneTree, moduleColors, main = "Network heatmap plot, all genes")
# dev.off()

# cat("18.2) Plotting network heatmap plot with selected genes.. \n")
# 
# nSelect = sel_genes
# # For reproducibility, we set the random seed
# set.seed(10);
# select = sample(nGenes, size = nSelect);
# selectTOM = dissTOM[select, select];
# # There's no simple way of restricting a clustering tree to a subset of genes, so we must re-cluster.
# selectTree = hclust(as.dist(selectTOM), method = "average")
# selectColors = moduleColors[select];
# # Open a graphical window
# # sizeGrWindow(9,9)
# # Taking the dissimilarity to a power, say 10, makes the plot more informative by effectively changing 
# # the color palette; setting the diagonal to NA also improves the clarity of the plot
# plotDiss = selectTOM^7;
# diag(plotDiss) = NA;
# pdf(file = paste(outputDir,"NetworkHeatmap_sel_genes.pdf", sep = '' ), width = 9, height = 9);
# TOMplot(plotDiss, selectTree, selectColors, main = "Network heatmap plot, selected genes")
# dev.off()

cat("19) Analyzing module eigengene networks.. \n")

# Recalculate module eigengenes
MEs = moduleEigengenes(datExpr, moduleColors)$eigengenes

# traitScores <- data.frame(matrix(ncol=length(names(datTraits)),nrow=nSamples, dimnames=list(NULL, names(datTraits))))
# for (trait in names(datTraits)){
#     traitScores[, trait] = datTraits[, trait]
# }

# Add the weight to existing module eigengenes
# MET = orderMEs(cbind(MEs, traitScores))
MET = orderMEs(cbind(MEs, datTraits))


# # Plot the relationships among the eigengenes and the trait
# # sizeGrWindow(5,7.5);
# pdf(file = paste(outputDir,"ME_dendrogram_heatmap.pdf", sep = '' ), width = 15, height = 7.5);
# par(cex = 0.9)
# plotEigengeneNetworks(MET, "", marDendro = c(0,4,1,2), marHeatmap = c(3,4,1,2), cex.lab = 0.8, xLabelsAngle
#                       = 90)

# Plot the dendrogram
# sizeGrWindow(6,6);
pdf(file = paste(outputDir,"ME_eigengene_dendrogram.pdf", sep = '' ), width = 10, height = 10);
par(cex = 1.0)
plotEigengeneNetworks(MET, "Eigengene dendrogram", marDendro = c(0,4,2,0), plotHeatmaps = FALSE)
dev.off()
pdf(file = paste(outputDir,"ME_eigengene_dendrogram_small.pdf", sep = '' ), width = 5, height = 3);
par(cex = 0.7)
plotEigengeneNetworks(MET, "Eigengene dendrogram", marDendro = c(0,4,2,0),colorLabels = T, 
                      plotHeatmaps = FALSE)
dev.off()
# Plot the heatmap matrix (note: this plot will overwrite the dendrogram plot)
pdf(file = paste(outputDir,"ME_heatmap.pdf", sep = '' ), width = 10, height = 10);
par(cex = 1.0)
plotEigengeneNetworks(MET, "Eigengene adjacency heatmap", 
                      heatmapColors = blueWhiteRed(50)[26:50],
                      marHeatmap = c(8,8,2,2), plotDendrograms = FALSE, xLabelsAngle = 90)
dev.off()

pdf(file = paste(outputDir,"ME_heatmap_filtered_blues.pdf", sep = '' ), width = 4.1, height = 3.5);
#par(cex = 1.0)
plotEigengeneNetworks(MET[,c("Age","Early", "Middle", "Late", "MEred","MEsalmon")], 
                      "Eigengene adjacency heatmap", 
                      marHeatmap = c(8,8,2,2), 
                      plotDendrograms = FALSE, 
                      # heatmapColors = brewer.pal(10, "Reds"),
                      heatmapColors = rev(blueWhiteRed(50)[1:25]),
                      xLabelsAngle = 90)
dev.off()

# #### create graphs
# 
# wgcna2igraph(bwnet, datExpr, top.n.edges = NA, modules2plot = "red",
#              colors2plot = NULL, kME.threshold = 0.75, adjacency.threshold = 0.1,
#              adj.power = 6, verbose = T, min.edge = 2, node.size = 0,
#              frame.color = NA, node.color = NA, edge.alpha = 0.5, edge.width = 1,
#              returnNet = TRUE)

##### GO analysis

library(clusterProfiler)
library(dplyr)
library(ggplot2)
library(org.Mm.eg.db)
library(enrichplot)

#####  

sel_modules <- c("salmon", "red", "brown", "darkorange", "black", "lightyellow", "tan",
                 "grey60", "darkgrey", "darkturquoise", "blue",
                 "purple", "skyblue", "white", "green", "yellow", "saddlebrown", "magenta")

gene_df <- geneInfo %>% filter(EnsemblIDs != "None")

list.clusters.go <- vector(mode="list", length=0)
for (i in sel_modules){
  modgeneInfo <- gene_df[gene_df$moduleColor == i,"GeneIDs"]
  list.clusters.go[i] <- list(i = modgeneInfo)
}

bp_go <- compareCluster(geneCluster = list.clusters.go, fun = "enrichGO", 
                        OrgDb='org.Mm.eg.db', keyType = 'SYMBOL',
                        ont = "BP", pvalueCutoff = 0.05, pAdjustMethod = "BH", 
                        qvalueCutoff = 0.1, universe = as.vector(geneInfo0$GeneIDs),
                        minGSSize = 10, maxGSSize = 500, readable = FALSE)

pdf(file = "/Users/methiaa/Downloads/GroupC_BulkRNA_WGCNA/output/GO_BP_selModules.pdf",
    width = 14, height = 13)
dotplot(bp_go, showCategory = 5) + 
  theme(axis.text.x = element_text(angle = 90, vjust = 0.5, hjust = 1, face = "bold"))
dev.off()

mf_go <- compareCluster(geneCluster = list.clusters.go, fun = "enrichGO", 
                        OrgDb='org.Mm.eg.db', keyType = 'SYMBOL',
                        ont = "MF", pvalueCutoff = 0.05, pAdjustMethod = "BH", 
                        qvalueCutoff = 0.1, universe = as.vector(geneInfo0$GeneIDs),
                        minGSSize = 10, maxGSSize = 500, readable = FALSE)

pdf(file = "/Users/methiaa/Downloads/GroupC_BulkRNA_WGCNA/output/GO_MF_selModules.pdf",
    width = 11, height = 11)
dotplot(mf_go, showCategory = 5) + 
  theme(axis.text.x = element_text(angle = 90, vjust = 0.5, hjust = 1, face = "bold"))
dev.off()

cc_go <- compareCluster(geneCluster = list.clusters.go, fun = "enrichGO", 
                        OrgDb='org.Mm.eg.db', keyType = 'SYMBOL',
                        ont = "CC", pvalueCutoff = 0.05, pAdjustMethod = "BH", 
                        qvalueCutoff = 0.1, universe = as.vector(geneInfo0$GeneIDs),
                        minGSSize = 10, maxGSSize = 500, readable = FALSE)

pdf(file = "/Users/methiaa/Downloads/GroupC_BulkRNA_WGCNA/output/GO_CC_selModules.pdf",
    width = 13, height = 13)
dotplot(cc_go, showCategory = 5) + 
  theme(axis.text.x = element_text(angle = 90, vjust = 0.5, hjust = 1, face = "bold"))
dev.off()

######## gene overlap analysis

library(GeneOverlap)
library(pheatmap)
library(Matrix)
library(RColorBrewer)

mod_df <- geneInfo %>% filter(EnsemblIDs != "None")
list.modGenes <- vector(mode="list", length=0)
for (i in unique(moduleColors)[-1]){
  modgeneInfo <- mod_df[mod_df$moduleColor == i,"GeneIDs"]
  list.modGenes[i] <- list(i = modgeneInfo)
}

cell.markers_df <- read.csv(file = "/Users/methiaa/Downloads/GroupC_BulkRNA_WGCNA/output/gene_overlap/top50_markers_per_type.csv")
list.cellGenes <- vector(mode="list", length=0)
for (i in names(cell.markers_df)[-1]){
  modgeneInfo <- as.data.frame(cell.markers_df[,c("gene",i)]) %>% filter(cell.markers_df[, i] != 0) %>% dplyr::select(gene)
  list.cellGenes[i] <- list(i = modgeneInfo$gene)
}

bgGenes <- geneInfo0$GeneIDs

gom.obj <- newGOM(gsetA = list.modGenes, gsetB = list.cellGenes, genome.size = length(bgGenes))

jacIdx_matrix <- as.data.frame(getMatrix(gom.obj, name="Jaccard"))
#jacIdx_matrix <- as.data.frame(getMatrix(gom.obj, name="odds.ratio"))
pval_matrix <- getMatrix(gom.obj, name="pval")
for(row in 1:nrow(pval_matrix)) {
  for(col in 1:ncol(pval_matrix)) {
    pval_matrix[row, col] <- as.numeric(format(round(pval_matrix[row, col], 2), nsmall = 2))
  }
}
pval_matrix[pval_matrix > 0.05] <- "N.S."

pval_x <- pval_matrix[apply(pval_matrix[,-1], 1, function(x) !all(x=="N.S.")),]
pval_x <- pval_x[apply(pval_x[-1,], 1, function(x) !all(x=="N.S.")),]
x <- jacIdx_matrix[row.names(pval_x), colnames(pval_x)]

go.p <- pheatmap(x,
                 show_rownames=TRUE,
                 show_colnames =TRUE,
                 cluster_cols=TRUE,
                 cluster_rows = TRUE,
                 treeheight_col = 0,
                 display_numbers = pval_x,
                 fontsize = 10,
                 fontsize_number = 5,
                 #number_format = "%.1f",
                 number_color = "black",
                 border_color = "black",
                 scale = "none",
                 color = colorRampPalette(brewer.pal(n = 9, name = "Reds"))(101),
                 #breaks = seq(0, 1, length.out = 101),
                 annotation_names_col = FALSE,
                 main = "Odds Ratio (Fischer's exact test)",
                 angle_col = 90)

# labeledHeatmap(Matrix = x,
#                xLabels = colnames(x),
#                yLabels = row.names(x),
#                yColorLabels = TRUE,
#                yLabelsPosition = "left",
#                colors = colorRampPalette(brewer.pal(9, "Reds"))(50),
#                textMatrix = pval_x,
#                setStdMargins = FALSE,
#                cex.text = 0.8,
#                zlim = c(0,0.1),
#                main = paste("Gene Overlap: Jaccard Index"))

png(filename = "/Users/methiaa/Downloads/GroupC_BulkRNA_WGCNA/output/gene_overlap/GO_celltype_wgcna.png",
    units = "in", width = 4.5, height = 2.9, res = 500)
go.p
dev.off()

gom.obj <- testGeneOverlap(gom.obj)

##

list.DGenes <- vector(mode="list", length=0)

DCA_df <- read.csv(file = "/Users/methiaa/Downloads/GroupC_BulkRNA_WGCNA/output/gene_overlap/initial_hubs.csv", sep = ";")
DCA_df <- data.frame(do.call('rbind', strsplit(as.character(DCA_df$genes),'.',fixed=TRUE)))
ngenes <- round(0.05*length(unique(DCA_df$X1)))
list.DGenes[1] <- list(DCA_hubs = unique(DCA_df$X1)[1:ngenes])
names(list.DGenes)[1] <- "DCA_hubs"

DEA_df <- read.csv(file = "/Users/methiaa/Downloads/GroupC_BulkRNA_WGCNA/output/gene_overlap/sig_diff_expression_gene_list.csv")
DEA_df <- data.frame(do.call('rbind', strsplit(as.character(DEA_df$X),'.',fixed=TRUE)))
list.DGenes[2] <- list(DEA_hubs = unique(DEA_df$X1))
names(list.DGenes)[2] <- "DEA_genes"

gom.obj <- newGOM(gsetA = list.modGenes, gsetB = list.DGenes, genome.size = length(bgGenes))

# jacIdx_matrix <- as.data.frame(getMatrix(gom.obj, name="Jaccard"))
jacIdx_matrix <- as.data.frame(getMatrix(gom.obj, name="odds.ratio"))
pval_matrix <- getMatrix(gom.obj, name="pval")
for(row in 1:nrow(pval_matrix)) {
  for(col in 1:ncol(pval_matrix)) {
    pval_matrix[row, col] <- as.numeric(format(round(pval_matrix[row, col], 2), nsmall = 2))
  }
}
pval_matrix[pval_matrix > 0.05] <- "N.S."

pval_x <- pval_matrix[apply(pval_matrix, 1, function(x) !all(x=="N.S.")),]
pval_x <- pval_x[apply(pval_x[-1,], 1, function(x) !all(x=="N.S.")),]
x <- jacIdx_matrix[row.names(pval_x), colnames(pval_x)]

go.p <- pheatmap(x,
                 show_rownames=TRUE,
                 show_colnames =TRUE,
                 cluster_cols=TRUE,
                 cluster_rows = TRUE,
                 treeheight_col = 0,
                 display_numbers = pval_x,
                 fontsize = 10,
                 fontsize_number = 5,
                 #number_format = "%.1f",
                 number_color = "black",
                 border_color = "black",
                 scale = "none",
                 color = colorRampPalette(brewer.pal(n = 9, name = "Reds"))(101),
                 #breaks = seq(0, 1, length.out = 101),
                 annotation_names_col = FALSE,
                 main = "Odds Ratio (Fischer's exact test)",
                 angle_col = 90)

png(filename = "/Users/methiaa/Downloads/GroupC_BulkRNA_WGCNA/output/gene_overlap/GO_Diff_wgcna.png",
    units = "in", width = 2.5, height = 2.5, res = 500)
go.p
dev.off()

# ##
# 
# DCA_df <- read.csv(file = "/Users/methiaa/Downloads/GroupC_BulkRNA_WGCNA/output/gene_overlap/initial_hubs.csv", sep = ";")
# DCA_df <- data.frame(do.call('rbind', strsplit(as.character(DCA_df$genes),'.',fixed=TRUE)))
# ngenes <- round(0.05*length(unique(DCA_df$X1)))
# list.DCAGenes <- list(DCA_hubs = unique(DCA_df$X1)[1:ngenes])
# 
# DEA_df <- read.csv(file = "/Users/methiaa/Downloads/GroupC_BulkRNA_WGCNA/output/gene_overlap/sig_diff_expression_gene_list.csv")
# DEA_df <- data.frame(do.call('rbind', strsplit(as.character(DEA_df$X),'.',fixed=TRUE)))
# list.DEAGenes <- list(DEA_hubs = unique(DEA_df$X1))
# 
# gom.obj <- newGOM(gsetA = list.DCAGenes, gsetB = list.DEAGenes, genome.size = length(bgGenes))
# 
# jacIdx_matrix <- as.data.frame(getMatrix(gom.obj, name="Jaccard"))
# pval_matrix <- getMatrix(gom.obj, name="pval")

### Eigengene boxplots

library("reshape2")
library("ggplot2")
library(ggpubr)
library(gtools)
library(tidyverse)
library("dplyr")
library("tidyr")

phenoData = read.csv("/Users/methiaa/Downloads/GroupC_BulkRNA_WGCNA//trait_data.csv", header = T);
print(dim(phenoData))
print(names(phenoData))
names(phenoData)[1] <- "Sample"

reqdSamples = rownames(datExpr);
traitRows = match(reqdSamples, phenoData$Sample);

datTraits = phenoData[traitRows, ];
rownames(datTraits) = phenoData[traitRows, 1];

MEs.df <- MEs
MEs.df$Type <- datTraits$Age_3bins

symnum.args <- list(cutpoints = c(0, 0.0001, 0.001, 0.01, 0.05, 1), symbols = c("****", "***", "**", "*", "ns"))

annot_col.me <- as.data.frame(datTraits[,3], row.names = datTraits[,1])
names(annot_col.me) <- "Age_groups"

MEs.df$Group <- annot_col.me$Age_groups

for (m in c("MEsalmon","MEred")){
  
  module <- substring(m, 3)
  mod_resultsDir <- paste(outputDir, "Age", "/", sep='')
  
  g <- ggplot(MEs.df, aes(x=Type, y=MEs.df[,m])) +
    geom_boxplot(outlier.shape=NA, aes(fill=Group)) + theme_classic() +
    theme(legend.position = "none") +
    theme_classic() + 
    scale_x_discrete(limits = c("Early", "Middle", "Late")) +
    theme(panel.grid.major = element_blank(), panel.grid.minor = element_blank(),
          axis.text.x = element_text(angle = 0)) +
    labs(
      title=paste("Module: ", module, "\n", sep = ""),
      x=NULL,
      y = "Module Eigenvalue") +
    stat_compare_means(symnum.args = NULL, 
                       mapping=aes(label = format.pval(..p.signif.., digits = 1)), 
                       comparisons = list(c("Early", "Middle"),
                                          c("Early", "Late"),
                                          c("Middle", "Late")), 
                       label.y.npc = 1,
                       method = "t.test")
  
  pdf(file = paste(mod_resultsDir, module, "_eigengene_boxplot_sig.pdf", sep = '' ), width = 5, height = 5);
  print(g)
  dev.off()
}
```
